# Exploring Single-Cell Data with Deep Multitasking Neural Networks

**DOI:** 10.1101/237065

**Authors:** Matthew Amodio, David van Dijk, Krishnan Srinivasan, William S Chen, Hussein Mohsen, Kevin R. Moon, Allison Campbell, Yujiao Zhao, Xiaomei Wang, Manjunatha Venkataswamy, Anita Desai, V. Ravi, Priti Kumar, Ruth Montgomery, Guy Wolf, Smita Krishnaswamy

## Abstract

Biomedical researchers are generating high-throughput, high-dimensional single-cell data at a staggering rate. As costs of data generation decrease, experimental design is moving towards measurement of many different single-cell samples in the same dataset. These samples can correspond to different patients, conditions, or treatments. While scalability of methods to datasets of these sizes is a challenge on its own, dealing with large-scale experimental design presents a whole new set of problems, including batch effects and sample comparison issues. Currently, there are no computational tools that can both handle large amounts of data in a scalable manner (many cells) and at the same time deal with many samples (many patients or conditions). Moreover, data analysis currently involves the use of different tools that each operate on their own data representation, not guaranteeing a synchronized analysis pipeline. For instance, data visualization methods can be disjoint and mismatched with the clustering method. For this purpose, we present SAUCIE, a deep neural network that leverages the high degree of parallelization and scalability offered by neural networks, as well as the deep representation of data that can be learned by them to perform many single-cell data analysis tasks, all on a unified representation.

A well-known limitation of neural networks is their interpretability. Our key contribution here are newly formulated regularizations (penalties) that render features learned in hidden layers of the neural network interpretable. When large multi-patient datasets are fed into SAUCIE, the various hidden layers contain denoised and batch-corrected data, a low dimensional visualization, unsupervised clustering, as well as other information that can be used to explore the data. We show this capability by analyzing a newly generated 180-sample dataset consisting of T cells from dengue patients in India, measured with mass cytometry. We show that SAUCIE, for the first time, can batch correct and process this 11-million cell data to identify cluster-based signatures of acute dengue infection and create a patient manifold, stratifying immune response to dengue on the basis of single-cell measurements.

## 1 Introduction

Vast amounts of high-dimensional, high-throughput, single-cell data measuring various aspects of cells including mRNA molecules, proteins, epigenetic marks and histone modifications are being generated via new technologies. Furthermore, the number of samples included in large-scale studies of single-cell data for comparing across populations or disease conditions is rapidly increasing. Processing data of this dimensionality and scale is an inherently difficult prospect, especially considering the degree of noise, batch effects, artifacts, sparsity and heterogeneity in the data [1,2]. However, this effect becomes exacerbated as one tries to compare between samples, which themselves contain noisy heterogeneous compositions of cellular populations.

Deep learning offers promise as a technique for handling the size and dimensionality of modern biological datasets. However, while work has been done in training networks to perform certain supervised tasks such as predicting binding [3,4] or classifying patients [5], deep learning has been underutilized for unsupervised exploratory tasks. In this paper, we develop a deep learning framework that focuses on unsupervised data exploration. Our key insight is that the layers of a deep neural network form representations of the data, and that if those layers are properly constrained (via architectural choices and regularization), they can be used to extract task-oriented features of the data.

We base our approach on the *autoencoder* [6–8]. An autoencoder is a neural network that learns to recreate its own input via a low-dimensional bottleneck layer that learns representations of the data and enables a denoised reconstruction of the input from them [9–13]. Since autoencoders learn their own features, they can reveal structure in the data without defining or explicitly learning a similarity or distance metric in the original data space as other dimensionality reduction methods do (for instance, PCA uses covariance and diffusion maps [14] utilize affinities based on a kernel choice). We use this approach to construct SAUCIE, a Sparse Autoencoder for Unsupervised Clustering, Imputation, and Embedding, which is aimed to enable exploratory tasks via its design choices.

SAUCIE is a multilayered deep neural network, whose input layer is fed single-cell measurements, such as mass cytometry or single-cell RNA sequencing, of an individual cell. Then, SAUCIE gradually reduces the dimensionality of the dataset by taking the data through narrower and narrower hidden layers. We see that the output or reconstruction layer of SAUCIE gives similarly denoised and imputed data as the manifold denoising method MAGIC [15] on a million single-cell RNA sequencing dataset from embryonic mouse brain. In other words, SAUCIE effectively learns the manifold of the data in a similar way to data diffusion [16] methods. Thus, SAUCIE can leverage the power of manifold learning, which has shown to be key for analyzing single-cell data [17] in a scalable fashion. Manifold learning methods are traditionally difficult to scale due to the computational complexity of kernel computation and eigendecomposition operations. Deep learning comes to the rescue here by being amenable to GPU speedup and parallelization of matrix operations.

As SAUCIE reduces input dimensionality, regularizations on different layers reveal different representations of the data: for visualization, batch correction, clustering, and denoising. In order to achieve these representations we use customized regularizations in each layer. We use the architectural choice of having a two-dimensional bottleneck layer to provide a *visualization* of the data. We develop a novel batch-level maximal mean discrepancy (MMD)-based penalty constraint to *remove batch effects* in the embedding layer. A customized sparse encoding layer featuring our novel information-dimension (ID) regularization provides an *automated clustering* of the data with no parametric assumptions on the shape or number of clusters. All regularizations balance against reconstruction accuracy, which is the basic penalty in an autoencoder that steers the network convergence away from trivial solutions. Furthermore, this penalty ensures that the final layer of the network provides reconstructed measurements that are *denoised*; in the case of single-cell RNA sequencing data, this layer also naturally *imputes missing values*.

Guiding the internal representations of the data to be effective at each of these disparate tasks together fit SAUCIE into the field of multitask learning. Results in multitask learning have generally shown that optimizing multiple tasks over the same latent representation is helpful in increasing the reliability and consistency of various algorithms. We apply the same approach here by having the representation (or data manifold) learned by SAUCIE be jointly optimized for multiple tasks. Further, SAUCIE itself forms a near complete analysis of the data. The clustering layer in SAUCIE for instance, actually performs clustering, and clusters are read out from this layer. This is in contrast to other methods that simply use the autoencoder for coming up with a reduced dimensional representation, which is then fed to other (generally unscalable) algorithms, for example scVI which outputs a latent layer that then needs another clustering algorithm [18].

We apply SAUCIE to a twenty-million cell mass cytometry dataset with 180 samples from forty subjects in a study of the dengue flavivirus [19]. SAUCIE is the only method that is able to batch correct 180 samples and then cluster them in such a way that subpopulation proportions become comparable prima facia. This obviates the need for approaches such as first clustering samples separately and then performing “meta-clustering” as with the Phenograph method, or other methods that cannot operate uniformly on combined data of this size (the problems of which are illustrated in Figure S10). We are also able to tune the granularity of clustering with SAUCIE in order to get a clustering that is informative of the differences between conditions. SAUCIE results show that acute subjects are characterized by enrichment in distinct subpopulations of CD4-CD8-*γδ* T cells and cells involved in Type I interferon signaling. When subjects are measured in convalescence, there is an increase in CD4+Foxp3+ T reg cells.

Thus, SAUCIE provides a unified representation of data where different aspects or features are emphasized in different layers, forming a one-step data analysis pipeline. This unified analysis uncovers a cell-space manifold as well as a sample-space manifold, thus enabling a multilevel analysis of complex experimental design where the samples are stratified on the basis of their cell-level features. We additionally evaluate SAUCIE extensively on all of its designed tasks using ten public single-cell datasets.

## 2 Results

### 2.1 The SAUCIE Architecture and Layer Regularizations

To enable unsupervised learning in a scalable manner, we base our method on the autoencoder. Autoencoders learn to recreate their input at the output layer, but via a low-dimensional informational bottleneck layers which are forced to learn meaningful structure-preserving representations of the data. However, a key challenge is to extract meaning from this representation. Specifically, we seek representations in hidden layers that are useful for performing the various analysis tasks associated with single cell data. Here, we introduce several design decisions and novel regularizations to our autoencoder architecture (Figure 1) in order to constrain the learned representations for four key tasks:

1. visualization and dimensionality reduction,
2. batch correction,
3. clustering, and
4. denoising and imputation.

**Figure 1:**
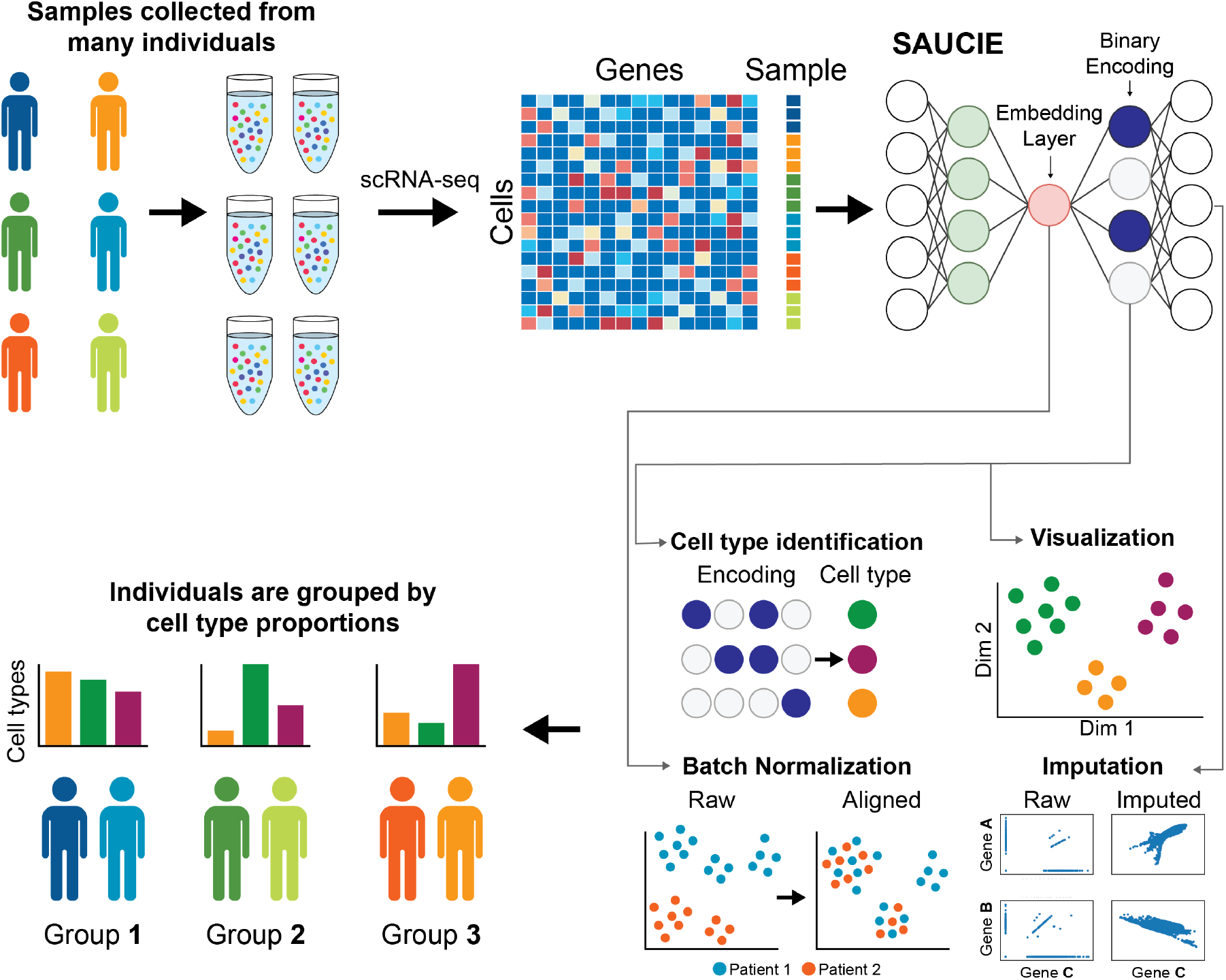
The pipeline for analyzing single-cell data in large cohorts with SAUCIE. Many individual patients are separately measured with a single-cell technology such as CyTOF or scRNA-seq, producing distinct datasets for each patient. SAUCIE performs imputation and denoising, batch effect removal, clustering, and visualization on the entire cohort with a unified model and is able to provide interpretable, quantifiable metrics on each subject or group of subjects.

For each task, dedicated design decisions are used to produce the desirable result.

#### Clustering

First, to cluster the data, we introduce the *information dimension* regularization that encourages activations of the neurons in a hidden layer of the network to be binarizable. The idea is that if we can obtain a “digital” binary encoding, then we can easily turn these codes into clusters. As Figure 2A shows, the network without regularizations tends to store its information in a distributed, or “analog” way. With the ID regularization the activations are all near 0 or 1, i.e., binary or “digital”, and thus amenable to clustering by simple thresholding-based binarization. As seen in Figure 3A, this leads to a clustering of the cells that effectively represents the data space. Thus, the ID regularization achieves an analog-to-digital conversion that enables interpretation of the representation as data groups or clusters corresponding to each binary code. A previous work in the same vein, Binary Connect, has shown the promise in encouraging networks to learn in ways that are easy to binarize. That work differs from SAUCIE though, in that they learn binary weights rather than binary activations, along with the goal being to improve computational efficiency rather than achieve a clustering of the data [20].

**Figure 2:**
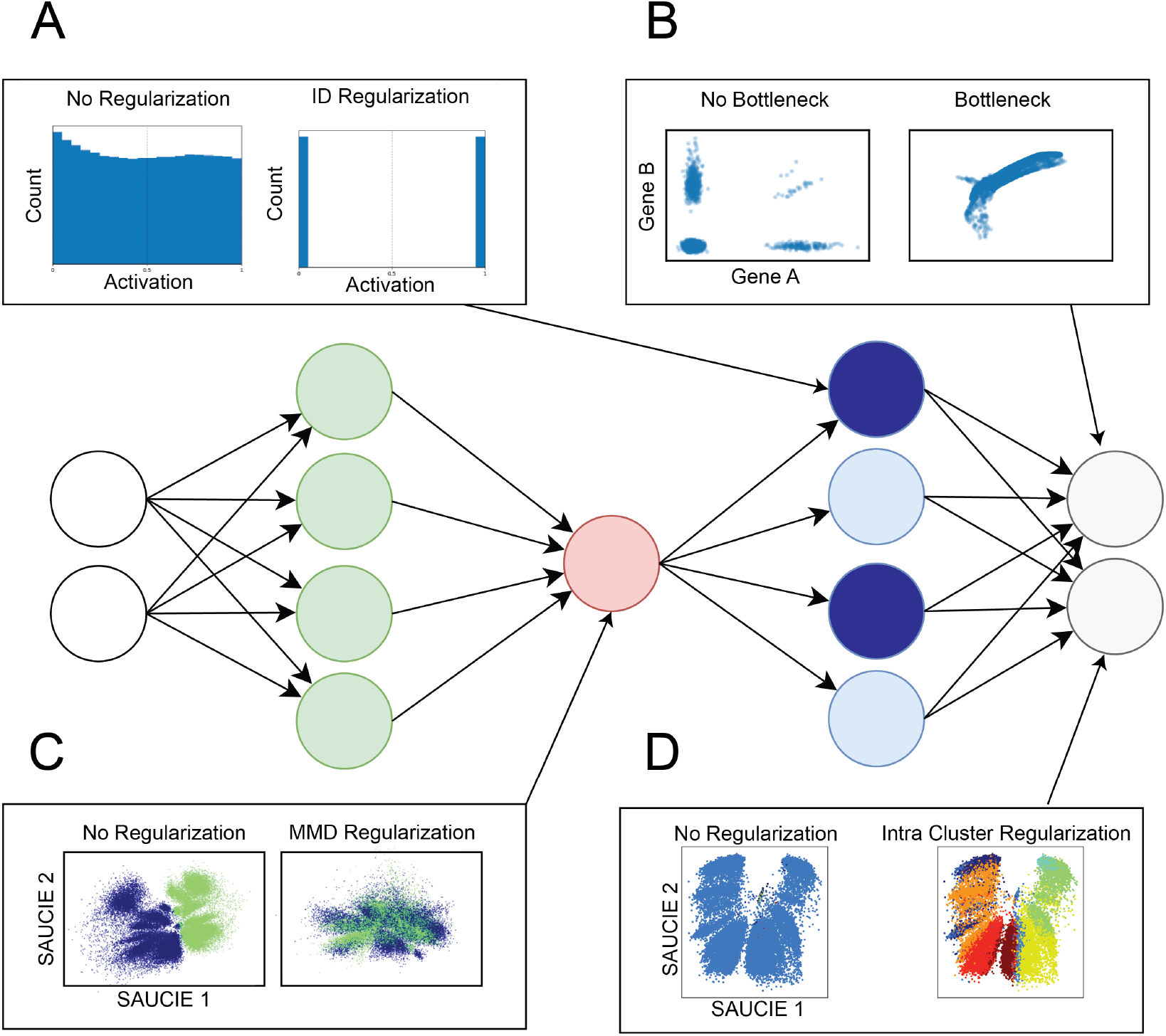
Regularizations and architecture choices in SAUCIE. A) the ID regularization applied on the sparse encoding layer produces digital codes for clustering B) the informational bottleneck, i.e. a smaller embedding layer, uses dimensionality reduction to produce denoised data at the output C) the MMD regularization removes batch artifacts D) the within cluster distance regularization applied to the denoised data provides coherent clusters.

**Figure 3:**
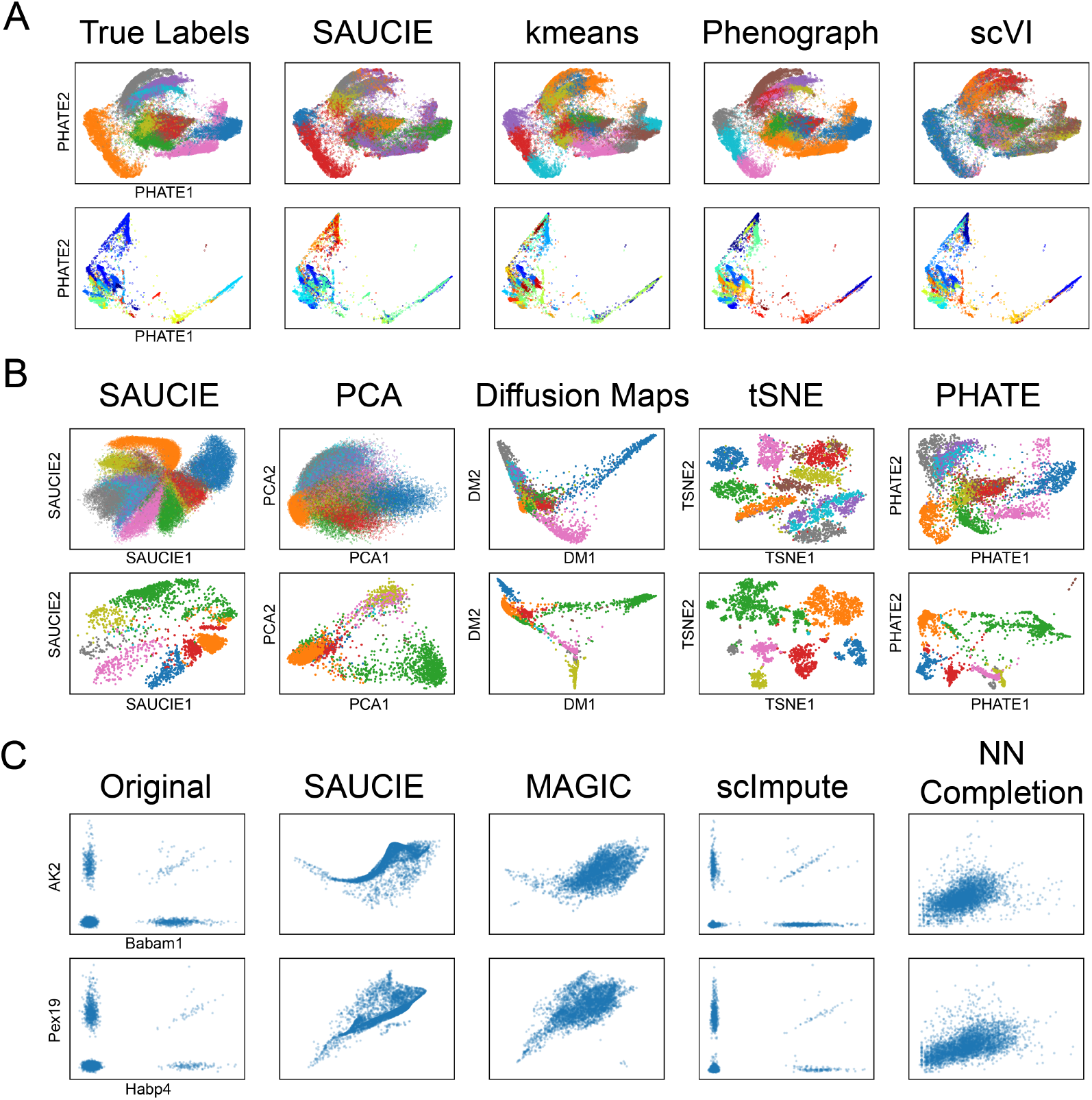
A comparison of the different analysis tasks performed by SAUCIE against other methods. A) A comparison of clustering performance shown on PHATE. SAUCIE compares well to the other methods, producing a coherent clustering. Neither Phenograph nor scVI produces clusters that look coherent. B) A comparison of SAUCIE’s visualization. PCA produces a blurry visualization. Diffusion maps shows a much simplified structure. tSNE shatters the space. SAUCIE produces a result similar to PHATE, revealing the structure in the data. C) A comparison of imputation. SAUCIE recovers complex nonlinear shapes of gene-gene relationships.

#### Batch Correction

Batch effects are generally systematic differences found in biological data measured under different experimental runs, largely due to ambient conditions such as temperature, machine calibration or day-to-day variation in measurement efficiency. Thus, measurements even from very similar systems, such as blood cells of the same patient, appear to have a shift or difference between two different experimental runs. To solve this problem, we introduce a maximal mean discrepancy (MMD) correction that penalizes differences between the probability distributions internal activations of samples. Previous work has attempted batch correction by minimizing MMD. However, those models assume that batch effects are minor and simple shifts close to the identity function, which is often the case [21]. Moreover, minimizing MMD alone only removes any and all differences between batches. In contrast, the additional autoencoder reconstruction penalty in SAUCIE forces it to preserve the original structure in each batch, balancing the goals of, on one hand, making the two batches alike while on the other hand not changing them. We note that this notion of a biological batch (data measured or run together) is distinct from the mini-batches used in stochastic gradient descent to train neural networks and the two should not be confused. The term batch is exclusively used to describe biological batches and when training with stochastic gradient descent the term mini-batches is used.

Figure 4 shows that analyzing data before batch correction can lead to misleading results, as artificial variation from batch effects can drown out the relevant variation within the biology that we are interested in. Penalizing MMD directly on the input space would be a flawed way of addressing batch effects because it would require making the assumption of (and thus being sensitive to the choice of) meaningful distance and similarity measures on the input points. Since the data is noisy and possibly sparse, by instead penalizing MMD on an internal layer of the network, we can correct complex, highly nonlinear batch effects by aligning points on a data manifold represented in these layers.

**Figure 4:**
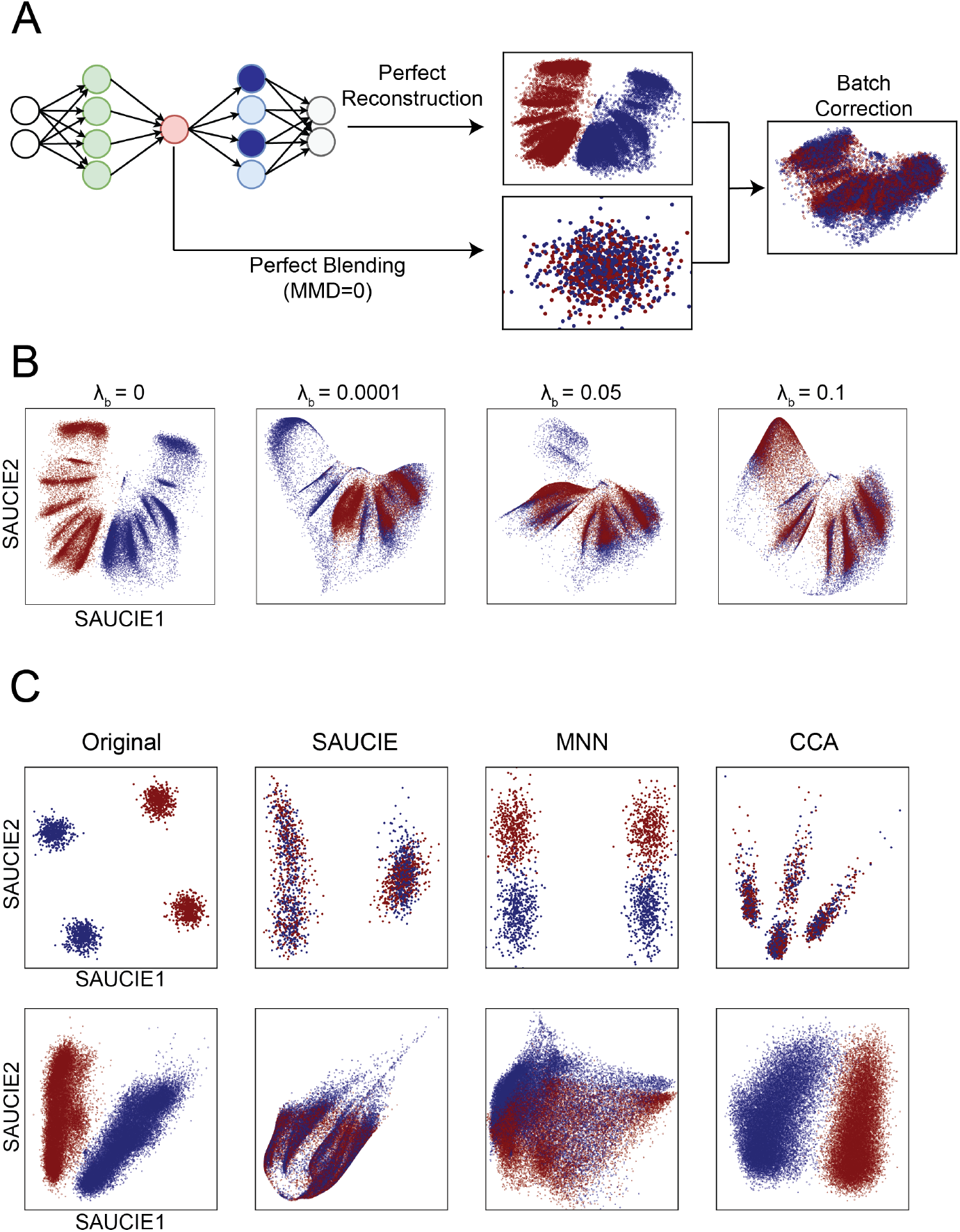
Demonstration of SAUCIE’s batch correction abilities. A) SAUCIE batch correction balances perfect reconstruction (which would leave the batches uncorrected) with perfect blending (which would remove all of the original structure in the data) to remove the technical variation while preserving the biological variation. B) The effect of increasing the magnitude of the MMD regularization on the dengue data. Sufficient MMD regularization is capable of fully removing batch effect. C) Results of batch correction on the synthetic GMM data (top) and the dengue data (bottom) shows that SAUCIE better removes batch effects than MNN and better preserves the structure of the data than CCA.

#### Imputation and denoising

Next, we leverage the fact that an autoencoder does not reconstruct its input exactly, but instead must learn a lower dimensional representation of the data, and decode this representation for data reconstruction. This means the reconstructions are denoised versions of the input and are thus naturally solutions to the dropout and other noise afflicting much real-world data, especially single cell RNA-sequencing data. The gene-gene relationships plotted in Figure 3C illustrate the ability of SAUCIE to recover the meaningful relationship between genes despite the noise in the data.

#### Visualization

Finally, we design the informational bottleneck layer of the autoencoder to be two dimensional, which lets it serve as a visualization and nonlinear embedding of the data. Because the network must reconstruct the input accurately from this internal representation, it must compress all the information about a cell into just these two dimensions, unlike methods like PCA or Diffusion Maps, which explicitly leave some variation unmodeled. Consequently, the information stored is also global, meaning points close together in the SAUCIE visualization are more similar than points that are farther apart, which is not true beyond small neighborhoods in a local method like tSNE. The ability to flexibly learn and accurately reflect the structure in the data with SAUCIE is demonstrated in Figure 3B.

Considered together, these customized regularizations and architectural choices make SAUCIE ideally suited for the exploratory data analysis when presented with single-cell biological data. Further, SAUCIE is entirely self-contained and not require any external algorithms that may not be able to process the scale of multisample single-cell data.

### 2.2 Comparison to other methods

We begin by offering an extensive comparison between SAUCIE and other (generally specialized) methods at each of these tasks in turn. We find that SAUCIE performs as well as, or even better than, specialized algorithms, which are much less scalable, for each individual task. Moreover, SAUCIE performs all tasks on a unified representation leading to visualizations that are coherent with clusters and cluster expression.

Throughout the comparisons on each of the tasks, we use two artificial datasets (simulation from mixtures of Gaussians and the canonical MNIST handwritten digit dataset), along with ten different single-cell datasets. Five datasets are CyTOF: the dengue dataset we extensively evaluate later in the manuscript, T cell development data from [22], renal cell carcinoma data from [23], breast tumor data from [24], and iPSC data from [25]. Five datasets are scRNA-seq: mouse cortex data, retinal bipolar cells from [26], hematopoiesis data from [27], mouse brain data from [28], and the 10x mouse megacell demonstration from [29].

#### 2.2.1 Clustering

To evaluate the ability of SAUCIE to find meaningful clusters in single-cell data, we compare it to several alternative methods: minibatch kmeans [30], Phenograph [31], and another neural network approach called Single-cell Variational Inference (scVI) [18]. While we compare to scVI as it and SAUCIE are both neural networks, we emphasize a fundamental difference between the two: scVI only returns a latent space, which must then be visualized or clustered by another outside method, while SAUCIE explicitly performs these tasks. Since kmeans needs to be told how many clusters there are ahead of time (k), we use the number of clusters identified by Phenograph as k. We look at the following datasets: MNIST handwritten digits for which there are ground truth labels, artificially generated Gaussians rotated into high dimensions, and public single-cell datasets for which we have curated cell clusters as presented by the authors: [26], [23], [28], [27], and [22].

In addition to analyzing the clusters visually (Figure S2), we also quantitatively assess cluster performance of the methods by computing modularity and silhouette scores [30] on the generated clusters and ground truth labels (Table 1). For MNIST, we find that just as we would expect given they are both non-Euclidean clustering methods that do not need a specified number of clusters, SAUCIE and Phenograph are the most comparable, with their having the highest modularities, similar silhouette scores, and very similar visual appearance. Next, we look at an artificially generated dataset of four two-dimensional Gaussian point clouds with different means rotated into 100 dimensions. We find that SAUCIE is the only method that automatically identifies exactly four clusters, which was the underlying number of clusters in the generation model. This illustrates why optimizing modularity, like Phenograph does, is not necessarily the best heuristic to follow, as it adds additional complexity to the clustering in order to increase the modularity score, resulting in too many clusters. Likewise, scVI did not identify the four clusters, which is unsurprising as the data did not fit its parametric model appropriate for gene counts.

**Table 1:**
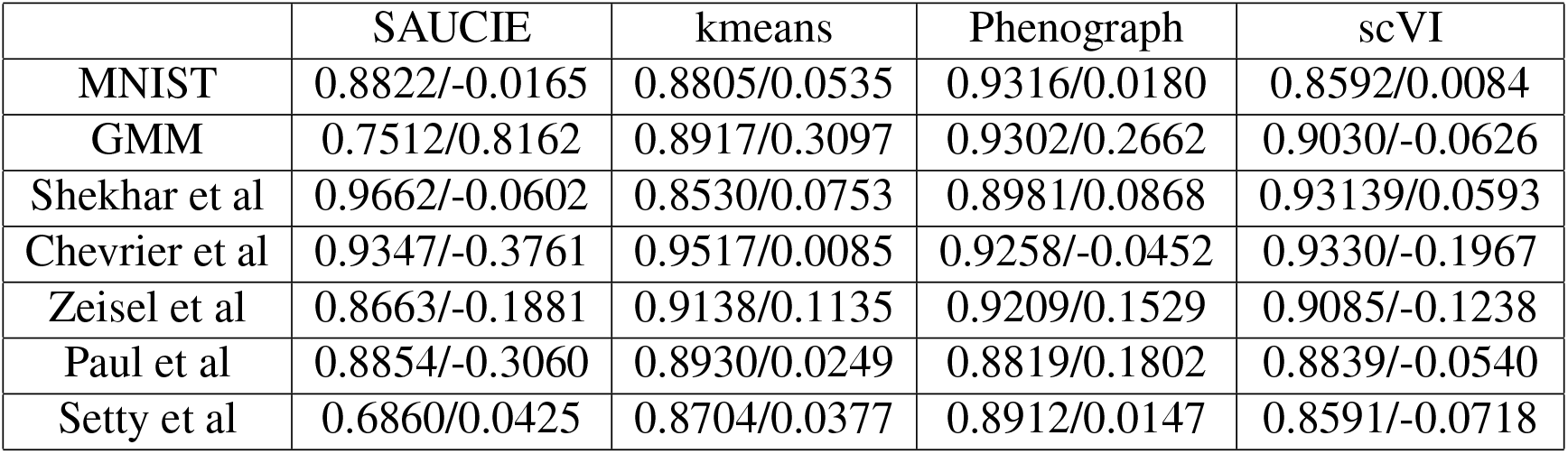
A comparison of modularity (left) and silhouette (right) scores of each of the clustering algorithms on each dataset.

We also examine clustering performance on five public single-cell datasets to evaluate the ability of SAUCIE to cluster real biological data: from [26], [23], [28], [27], and [22]. Visual inspection reveals that SAUCIE produces clusters that are qualitatively coherent on the embedding. Quantitatively, the modularity scores of its clusters corroborate this evaluation. As shown in Table 1 the average modularity score across datasets is 0.8531. In a wide variety of data from both CyTOF and scRNA-seq measurements, SAUCIE is able to produce clusters that reasonably represent the data qualitatively, quantitatively, and by comparison to other methods.

#### 2.2.2 Batch correction

We assess our ability to remove batch-related artifacts with SAUCIE by comparison to two published batch correction methods that have been specifically designed to remove batch effects in single-cell data. The first, Mutual Nearest Neighbors (MNN) [32], uses mutual nearest neighbors on a k-nearest neighbors graph to align two datasets, and the second, Canonical Correlation Analysis [33], finds a latent space in which the two batches are aligned. To evaluate the performance of these methods and SAUCIE, we use several different datasets with varying degrees of batch artifacts. We note that SAUCIE is the only method capable of scaling batch correction to hundreds of samples as we do in the next section. Nonetheless, here we compare performance on datasets small enough for the alternative methods to handle.

To quantitatively assess the quality, we apply a test we term the *mixing score* (similar to that of [34]):

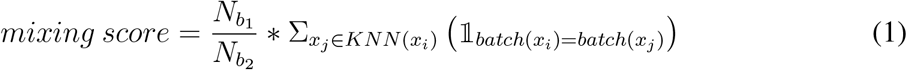

where *N*_*b*_1__ and *N*_*b*_2__ are the number of points in the first and second batch respectively. This score calculates for each point the number of nearest neighbors that are in the same batch as that point, accounting for the difference in batch sizes. In perfectly mixed batches, this score is 0.5, while in perfectly separated batches it is 1.0. As batch correction should not only mix the batches but also preserve their shape as best as possible, we quantify the distortion between the original and batch corrected data using Procrustes, which finds the error between the optimal alignments of the two batches by linear transformation [35]. These numbers are reported in Table 2. While the other methods each have some datasets that violate their assumptions and thus they perform poorly, SAUCIE performs as well or better at each of the wide variety of datasets.

**Table 2:** A comparison of mixing (left) and Procrustes (right) scores of each of the batch correction algorithms on each dataset.

|  | Original | SAUCIE | MNN | CCA |
| --- | --- | --- | --- | --- |
| GMM | 0.999/— | 0.630/0.629 | 0.526/0.620 | 0.510/0.998 |
| Dengue | 0.999/— | 0.593/0.532 | 0.998/0.512 | 0.992/0.765 |
| Mouse cortex | 0.994/— | 0.530/0.498 | 0.898/0.485 | 0.836/0.923 |
| Chevrier et al | 0.880/— | 0.540/0.934 | 0.787/0.232 | 0.835/0.346 |
| Azizi et al | 0.621/— | 0.512/0.180 | 0.560/0.205 | 0.621/0.000 |
| Setty et al | 0.518/— | 0.504/0.064 | 0.514/0.067 | 0.523/0.698 |

First, we generated two batches, each consisting of two ten-dimensional Gaussian point clouds with different means. We then rotated this into 1000 dimensions to simulate realistic single-cell data. Visual inspection shows that CCA appears to align the batches (i.e., the batch label is well mixed), however it distorts the original shape of the data, creating more distinct clusters per batch than originally existed. MNN pulls the batches closer together but does not fully mix them. SAUCIE appears to successfully align the two batches while at the same time preserving the original data structure shape without distortion. SAUCIE scores as well as the alternative methods at the mixing score, while only SAUCIE can easily scale this performance to hundreds of batches.

Next, we look at the CyTOF measurements of spike-in data where the same blood sample has been measured twice on different days. Since they are technical replicates, the difference between them confirms that there are batch effects in this data that need to be corrected. We expect perfect alignment after batch correction. We can observe well-aligned batches for SAUCIE and MNN, however CCA does not remove any batch effect. As before, SAUCIE scores well both in the mixing score and the Procrustes score.

Then, we evaluate nontechnical replicates of scRNA-seq data from developing mouse cortex. While the batch effect is the dominant signal in the data, we do not expect perfect alignment, as there are also possible differences between the time points that we expect to remain (the two samples are from embryonic day 14.5 and 17, respectively). CCA partially aligns the two batches. However, batch effect remains the strongest signal in the embedding and the shape of the data has been distorted: there now appear to be more clusters than were present originally in the data. SAUCIE and MNN, however, well align the two batches, but like in previous datasets, MNN appears to also remove much of the population structure of the data. SAUCIE both preserves the original population structure of each sample and aligns them. This is also reflected in the nearest neighbor values, which are 0.544, 0.689, and 0.902, respectively.

SAUCIE is also able to correct varying degrees of batch effect on public datasets from [23], [24], and [22]. With an average mixing score of 0.518 across the datasets, SAUCIE effectively aligns each different pair appropriately, due to its combination of reconstruction penalty and batch correction term. Both on scRNA-seq and CyTOF data, SAUCIE can integrate different samples for later downstream analysis.

#### 2.2.3 Visualization

To evaluate the SAUCIE visualization and its ability to provide a faithful low-dimensional data representation, we provide an extensive comparisons of this visualization to other frequently used methods. We make use of artificial datasets where the underlying structure is known, as well as real biological datasets that have been extensively characterized previously, so we have prior understanding of the structure we expect to see in the visualization (Figure S4).

The first three datasets come from a continuous artificially-generated tree structure with different amounts of Gaussian noise added to it. All seven of the branches are recovered by SAUCIE, tSNE, and PHATE. However, without enough noise, tSNE shatters branches, misleadingly showing them as different clusters. PCA, Monocle2, and Diffusion Maps correctly display the continuous tree-like nature of the data. However, in the two dimensions that are shown, they do not capture all of the branches.

In the tree generated using diffusion limited aggregation (DLA), we have a more complicated tree than in the previous examples. Only SAUCIE and PHATE effectively illustrate this branching structure, while PCA places spherical clouds with many branches overlapping, and Monocle2 and Diffusion Maps collapse several of the branches together. tSNE shatters the different branches into one or more clusters, losing the continuous nature.

Next, to evaluate the ability of the various embedding methods to handle intersecting manifolds, we generated a dataset of three intersecting half circles. Both SAUCIE and PCA preserve the circular shape as well as the intersecting positions. The other methods either distort the curvature of the data, shatter the trajectory, or remove the intersecting nature of the data.

To evaluate the ability of SAUCIE and the existing visualization methods to recover underlying structure we embed the MNIST dataset where there are true labels that correspond to the digit each image represents. We find that these different digits are well represented by SAUCIE, tSNE, and PHATE. In PCA, Monocle2, and Diffusion Maps, only some of the digits are distinct in the two dimensions that are shown, with the others being erroneously blended.

Another dataset where we have ground truth is a synthetic Gaussian mixture model (GMM). Here, four shifted Gaussians represented in the GMM dataset show the ability of each method to capture the distinct clusters present in the data. Diffusion Maps collapses all of the data into a single point in the two dimensions shown, while Monocle2 places the clusters closer or farther to each other erroneously. Additionally, PCA, Monocle2, and Diffusion Maps do not capture the spherical structure of the data. SAUCIE, tSNE, and PHATE all capture this structure effectively.

In [27], the authors performed an extensive characterization of hematopoiesis in mouse bone marrow and identified different cell types as shown in the colors in the embedding. SAUCIE produces a visualization that reflects branching structure that is consistent with PHATE. Monocle2 and Diffusion Maps collapse the trajectories into a single branch while tSNE shows them as contiguous clusters.

The data from [22] describes a system of T cell development in the mouse thymus in which T cells develop from CD4-CD8 double negative phenotype into double positive and then branch out into CD4+/CD8- and CD4-/CD8+. We therefore expect the embedding to show a continuous trajectory that then branches into two. This is the case for SAUCIE and PHATE. While tSNE shows the two directions, it does not optimally show the continuous progression. PCA and Monocle2 show a continuous progression but fail to show the branch point. Diffusion Maps fails to accurately capture any meaningful structure at all.

Next we looked at the dataset of [25] with induced pluripotent stem cells that were measured in CyTOF over the course of several days, denoted by different colors. We expect the time points to correlate with the embedding as cells gradually change phenotype over time. We can see that SAUCIE, PHATE, tSNE, and Diffusion Maps show this significant separation. PCA and Monocle2 show the least separation across time.

In [26], we examine retinal bipolar cells, along with the different subtypes identified by the authors. We expect the embedding to reflect these different populations that they identified. We can see that PHATE, tSNE, and SAUCIE are able to show all of the different clusters within the two dimensional embedding. PCA, Monocle2, and Diffusion Maps show some of the structure but clearly do not show all of the distinctions between cell types.

In [28], we look at mouse neural cells, which were also accompanied by different neural cell types that are reflected by different colors in the embeddings. Again we find that SAUCIE, PHATE, and tSNE show all the expected cell types and that PCA, Monocle2, and Diffusion Maps only capture some of the structure within the two dimensions that are shown.

In addition to the previous extensive qualitative evaluation, we also measure the quality of the visualizations with a quantitative metric taken from [36]. In line with to their method’s precision and recall metrics, we compute a neighborhood around each point in both the original data space and the embedding space, and compare the neighbors of each. An embedding with high recall has most of a point’s original-space neighbors in its embedding-space neighborhood. Similarly, an embedding with high precision has most of the point’s embedding-space neighbors in its original-space neighborhood. As directed by the authors’ algorithm, we gradually increase the size of the neighborhood and report the area-under-the-curve (AUC) for the precision-recall curve. These results are in Table 3, where SAUCIE has the highest average score of 0.9342, averaged across all datasets.

**Table 3:** A comparison of precision-recall area-under-the-curves (AUCs) for each of the visualization algorithms on each dataset.

|  | SAUCIE | PCA | Monocle2 | DM | UMAP | TSNE | PHATE |
| --- | --- | --- | --- | --- | --- | --- | --- |
| Artificial Tree 3 | 0.993 | 0.956 | 0.935 | 0.963 | 0.967 | 0.968 | 0.947 |
| Artificial Tree 7 | 0.994 | 0.951 | 0.921 | 0.980 | 0.986 | 0.990 | 0.971 |
| Artificial Tree 20 | 0.948 | 0.896 | 0.854 | 0.940 | 0.938 | 0.940 | 0.939 |
| DLA Tree | 0.865 | 0.817 | 0.725 | 0.819 | 0.836 | 0.845 | 0.847 |
| Half Circles | 0.975 | 0.970 | 0.940 | 0.937 | 0.958 | 0.946 | 0.925 |
| GMM | 0.999 | 0.972 | 0.953 | 0.500 | 0.992 | 0.992 | 0.969 |
| MNIST | 0.777 | 0.801 | 0.744 | 0.498 | 0.718 | 0.506 | 0.507 |
| Paul et al | 0.944 | 0.948 | 0.896 | 0.807 | 0.842 | 0.865 | 0.856 |
| Setty et al | 0.882 | 0.870 | 0.839 | 0.508 | 0.501 | 0.501 | 0.491 |
| Zunder et al | 0.939 | 0.903 | 0.884 | 0.505 | 0.522 | 0.513 | 0.510 |
| Shekhar et al | 0.942 | 0.908 | 0.918 | 0.506 | 0.863 | 0.508 | 0.496 |
| Ziesel et al | 0.952 | 0.914 | 0.903 | 0.943 | 0.909 | 0.881 | 0.905 |

#### 2.2.4 Imputation

We analyze the SAUCIE imputation and its ability to recover missing values by implicitly interpolating on a data manifold in several ways. First, Figure S5 shows several relationships from the scRNA-seq data of the 10x mouse megacell dataset affected by severe dropout. This dataset consists of 1.3 million cells, and SAUCIE was the only method in the comparison to be able process the full dataset. Moreover, it was able to do this in just 44 minutes. Additionally, because training a neural network only requires small minibatches in memory at one time, we were able to do this without ever loading the entire large dataset into memory all at once. Thus, to enable this comparison, we subsampled the data by taking one of the SAUCIE clusters consisting of 4172 cells.

For this comparison, we measure against several popular imputation methods for scRNA-seq data: MAGIC, which is a data diffusion based approach, scImpute, which is a parametric statistical method for imputing dropouts in scRNA-seq data, and Nearest Neighbors Completion (NN Completion), which is an established method for filling in missing values in a general application of high-dimensional data processing.

In Figure S5, we show six relationships of the mouse megacell dataset for the original data and the different imputation methods. We observe that the original raw data is highly sparse, which can be seen by the large number of values on the axes where one of the variables is exactly zero. Note that most cells have one or both genes missing. This is a problem because this prevents us from identifying trends that exist between the genes. After imputation with SAUCIE, we can observe that the sparse character of the data has been removed, with values filled in that reveal underlying associations between the gene pairs. These associations are corroborated by MAGIC, which imputes similar values to SAUCIE in each case. MAGIC is a dedicated imputation tool that is widely used, so SAUCIE matching the relationships it found gives confidence in the ability of SAUCIE to impute dropout effectively. The resulting imputation in scImpute does not look significantly less sparse from the original and we do not see continuous trends emerge. NN Completion appears to desparsify the data, but the resulting trends all look similar to each other (i.e., positively correlated). This suggests that it does not correctly identify the underyling trends, as we would expect different genes to have different relationships. While scRNA-seq is highly sparse, the undersampling affects all entries in the matrix, including the nonzero values. As such, manifold-based methods like SAUCIE and MAGIC are more suited for finding these true relationships because they denoise the full dataset as opposed to just filling in zeros.

Due to the fact that ground truth values for the missing counts in this single-cell data are not known, we further test the accuracy of the imputation abilities of SAUCIE with an artificially constructed experiment. We first leverage the bulk RNA sequencing data of 1076 cells from [37], because it accurately captures the relationships between genes due to it not being sparse (as opposed to generating our own synthetic data from a parametric generating function that we have the ability to choose, where we can create the relationships). We then simulate increasing amounts of dropout and compare the imputed values returned by each method to the true values we started with. To simulate dropout in a manner that reflects the underlying mechanisms of inefficient mRNA capture, we remove *molecules* instead of just setting values for genes to zero. As a result, the level of dropout is conditional upon expression level, reflecting the dropout structure of single-cell RNA sequencing data. The results are reported in Figure S6, where SAUCIE compares favorably to other methods, recovering the true values accurately even after as much as 99% dropout. The dataset for this experiment consisted of just 1076 cells, which allowed us to compare to the methods that cannot process larger datasets, but even on a dataset of this size SAUCIE gave a more than 100-times speedup over NN Completion and 600-times speedup over scImpute.

#### 2.2.5 Runtime Comparison

In order to showcase the scalability of SAUCIE, we compare to a host of other methods on a subset of our newly generated CyTOF dataset consisting of over 11 million cells existing in 35 dimensions. We display the runtimes of each method on a random sample of *N* points, with *N* = 100, 200, 400, 800, …, 11000000 in Figure S1. For each step, the method was given a timeout after 24 hours. Points where a method stopped scaling in Figure S1 are marked with an ‘x’.

SAUCIE performs visualization, batch correction, imputation, and clustering in its run, while each of the other methods only performs one of these tasks. Moreover, SAUCIE does not just compute simple linear functions on the data, but instead performs complex non-linear transformations in the process. Despite its complexity, it also scales very well with the extremely large dataset sizes, which can be further improved by simply adding more independent GPUs for calculations. Each additional (relatively inexpensive) GPU can offer a near linear increase in computation time, as opposed to more CPUs which offer diminishing returns in parallelizability. All experiments were run on a single machine with just one GPU, meaning these results could still benefit even more from this potential for scalability. For further details on how the runtime experiment was performed, see the Methods section.

Among the batch correction methods, there are no other methods that correct multiple batches simultaneously. However even when we restrict to pairwise comparisons, SAUCIE is the only method that comes close to handling this amount of data. CCA and MNN both stop scaling in the tens of thousands of cells. In the group of imputation methods, scImpute and NN completion also stop scaling in the tens of thousands, while MAGIC stops scaling in the hundreds of thousands. For visualization, PCA was the only method faster than SAUCIE, which is unsurprising because calculating it using fast randomized SVD is quick, but it gives a simple, strictly linear blurry views of the data, in contrast to SAUCIE’s nonlinear dimensionality reduction. The other more complex visualization methods do not scale to these dataset sizes: Diffusion Maps, PHATE, tSNE, and Monocle2 all stop scaling before even reaching the full eleven million cells. For clustering, kmeans is the only one faster than SAUCIE, due to using its minibatched version. However, it still assumes circular clusters in the Euclidean space and comes with the intrinsic flaw that the number of clusters must be known ahead of time, which is not possible in any realistic setting like ours where we are performing exploratory data analysis on a large new dataset. Phenograph and scVI do not scale to the full dataset, either. Despite being another neural network method, scVI cannot scale to these larger sizes because it only produces a latent space that then must be clustered with another method. This requirement then becomes its bottleneck, emphasizing the importance of SAUCIE performing all tasks directly instead of acting as a pre-processing step for other methods.

SAUCIE is the only method that can efficiently batch correct, impute and denoise, visualize, and cluster datasets of this size, while using a nonlinear manifold representation of the data.

### 2.3 Analysis of immune response to dengue infection with SAUCIE

Next, we demonstrate an application of SAUCIE as an important tool enabling exploratory analysis of a new “big” dataset that consists of single-cell CyTOF measurements of T cells from 45 subjects including a group acutely infected with the dengue virus and healthy controls from the same endemic area [19]. While dengue is estimated to affect sixty million people yearly and cause ten thousand deaths, like other tropical diseases, it remains understudied. Moreover, dengue is especially challenging since there are several different serotypes with complex interactions between them. Specifically, there are four strains that have very different characteristics. While infection with a particular strain may provide some immunity towards reinfection with that same strain, an antibody dependent enhancement results in faster uptake of another strain upon reinfection [38]. Drugs have proven difficult to develop for dengue. Further, vaccine development has also been challenging in the case of dengue. Recently, the WHO has ruled that the dengue vaccine of Senofi Pasteur only be administered to patients who are infected for the second (or subsequent) time [39]. This is because the vaccine itself is thought to leave patients vulnerable to very severe reinfections. So unlike other viruses, the dengue virus apparently leaves patients more vulnerable the second time. These types of complex effects require deep and detailed analysis of both infected and convalescent patients at the single cell level to understand the immune response.

We applied SAUCIE to the single-cell CyTOF data of T cells collected in an area endemic for dengue virus infection [19] to study general T cell compartment composition, variability and changes in the variability after convalescence. We believe that the dengue data is an ideal test case for SAUCIE, because the samples are shipped from India and samples were collected over a period of months and were assesed over different experiment days [19]. Thus, there is a pressing need for batch correction and data cleaning as well as uniform processing, clustering and meta-analysis of patient stratification. As part of the study, cells from additional patient groups beyond the acutely infected were also measured: healthy people unrelated to the subjects as a control and the same acute subjects at a later convalescent time point. Primary research questions include understanding profile of the acute subjects and how they differ from the other groups. Across all groups, there are 180 samples resulting in over twenty million cells with results analyzed on 35 different protein markers, a massive amount of data that would cause difficulties in most standard analytic frameworks.

#### 2.3.1 Batch correction

Beyond the sheer size of the total dataset, due to the large number of distinct samples in the experiment there are significant batch related artifacts effects, stemming from day-to-day differences, instruments, handling and shipping of the samples. While there are true biological differences between the individual samples, to identify those true differences in the samples we have to remove differences that are caused by these technical variables.

Differences that are highly associated with the day they were run on the cytometry instrument can be seen by grouping all of the samples together by run day and examining their marker-by-marker abundances. Each run day has twelve samples chosen such that each day has samples from each experimental condition, so any differences between the samples from each day are batch effects. As shown in Figure S3, these difference exist in the spike-in controls as well as the samples, confirming their identity as batch effect and not true variation.

Figure S7 shows four markers with extreme batch effects: TCRgd, IL-6, IFNg, and CD86. These batch effects would normally mean only samples within each run day could be compared to each other, as comparisons between samples from different run days would be dominated by the differences in the run days. Instead, the SAUCIE batch correction removes these undesirable effects by combining the samples from each day and aligning them to a reference batch, here chosen to be Day 1. Figure S7 shows that after SAUCIE the differences between run days disappear so that now what it means to be low or high in a marker is the same for each day. Before, the cells with the lowest IFNg in samples from Day 3 would still be considered IFNg+ while the cells with the highest IFNg in samples from Day 1 would still be IFNg-. After batch correction with SAUCIE, these can be directly compared.

The challenge of batch correction is to remove differences due to artifacts while preserving biological differences. We reason that to prevent removing true biological variation, the ‘shape’ of the data (but not its position and scale) within each day must be preserved. We define the shape of the data as any moment beyond the first two - mean and variance. We examine this in detail by considering a run day with the most significant batch effects, Day 2. In Figure S3C, the SAUCIE visualization shows that the reference and nonreference batches are completely separated. When MMD regularization is added in SAUCIE, though, these two batches are fully overlapped. In Figure S8, we examine the twelve individual samples that were run on Day 2. Initially, we see that this confirms our idea that the differences between days are batch effects, because each sample measures high in IL-6 and CD86. So the differences between samples run on Day 1 and Day 2 in CD86 abundance is not dominated by having more of a certain sample type in Day 2. Instead, all samples in Day 2 have been shifted higher. As desired, after batch correction, the mean of each marker is reduced to the level of the reference-batch mean. Crucially, the relationship of samples in Day 2 relative to each other is preserved. The samples with the highest IL-6 in Day 2 are still Samples 3, 9, and 11 while the samples with the lowest are still Samples 4, 5, and 6. SAUCIE has just changed what it means to be high or low for samples in this day such that it reconciles what it means to be high or low for samples in the reference day.

In conclusion, the batch correction and denoising ability of SAUCIE has transformed the data into a form that is amenable to biological discovery. We investigate this in the next section.

#### 2.3.2 Differential cluster proportions between subjects

We first obtain the clusters characteristic of each group and then further analyze them for marker enrichments as single cell versions of blood biomarkers [40]. For the clustering considered here, we use a coarse-grained clustering obtained with a coefficient for ID regularization of 0.1. This was chosen by scanning across values of 0.01, 0.1, 0.2, 0.3, 0.4, and 0.5, and choosing the clustering that yielded the best modularity. If other granularities are desired, lower coefficients could be used and the impact of this parameter on the number of clusters is shown in Figure S9. The two regularizations *λ_d_* and *λ*_*c*_ affect the number of clusters that result. For a given value of *λ*_*d*_, as *λ*_*c*_ increases, the number of clusters decreases (coarser granularity). Higher values of *λ*_*d*_ yield more clusters (finer granularity). Notably, these results are robust and yield reasonable results for varying values of the two regularizations. These two together act as knobs that can be tuned to get the desired granularity of clustering. The methods section further discusses how these regularizations affect the number of clusters.

For the SAUCIE clustering, we focus on T cells as particularly relevant to the immune process and an abundant subset of the data (eleven million total cells), looking for clusters that are over- or under-represented in the cells of each group. We look for clusters that behave differently in the acute compared to the convalescent time points. These would then represent a population of cells that might have an important role in the process, which could be further investigated. To understand what cell population this is, we examine the marker abundance profile for the cluster. The mean for each cluster and marker is shown in the heatmap in Figure 6B.

We find twenty total clusters within the T cell populations, five of which are CD8 T cells and thirteen of which are CD4 T cells. In addition, interestingly, there are six clusters of CD4- CD8- T cells, where four are *γδ* T cells. These have been noted as a characteristic of reaction to viral infections [41–45]. There are twelve clusters representing effector memory cells and nine regulatory T cells that are CD4+Foxp3+. Two of the clusters are naive T cells.

Several of these populations are indicative of differences between acute, convalescent, and healthy subjects, and can be used for characterizing the nature of the reaction of each of these groups, as we do below.

1. *γδ* T cells are a relatively rare type of T cells, but SAUCIE is still able to identify them. Despite their rarity, they appear to have significance in identifying different populations, which emphasizes the importance of this attribute of SAUCIE. These cells signal especially strong earliy in immune response, particularly skin and mucosal immunity. They have less variable TCR sequences than *αβ* T cells [46]. These cells are a bridge between T cells and myeloid cells, as they have some innate immune activity, where they express CD11c and CD86. They can bind to lipid antigens. Clusters 0 and 3 (consisting of 7% of the total cells) shows upregulation of CD57. This is an indication of terminal differentiation. CTLA-4 and CD38 are also high, so these are highly activated cells and potentially dysfunctional. We see that these clusters are highest in the acute subjects and lowest in the healthy subjects. Out of the fifteen subjects that were measured both as acute subjects and later in convalescence, thirteen had more of these cells during their acute infection.
2. We find another group of *γδ* T cells that are CD45RO and CD45RA positive (cluster 2, consisting of 1% of the total cells), but not yet fully terminally differentiated, so these could be transitional between naïve and effector memory. The effector memory cells express less IFNb. As this cluster is more expressed in the healthy subjects, it indicates that even these subjects may have had some exposure to dengue. There is a lack of an inflammatory state, i.e., low in IFNb and Perforin, so we expect that these are actually memory cells instead of effector cells. It makes more sense then that these populations are more expressed in convalescent and healthy subjects.
3. We also find another population of CD4+ T cells (clusters 3-15, consisting of 45% of the total population) that are not expressing any inflammatory markers or activation markers, and these are higher in the convalescent and healthy subjects, while being very low in the acute subjects. These look to be other memory cells that may characterize these convalescent subjects. In fact, out of the fifteen subjects with acute-convalescent paired measurements, eleven had more of these cells during convalescent measurement. These have signs of recent activation as they do not have CD69, which is an early activation marker, nor any of the cytokines like IFNg, IFNb, or IL-6.
4. Additionally, we find a population of CD8+ effector cells (cluster 15, which consists of 3% of the total cells) that are highly expressed in the acute subjects. These cells also express CD57 and CD38, but are not *γδ* as the previous populations were. These appear to be more differentiated and are likely not transitional, as the previous ones were, either.

We can also visualize the cell-level cluster proportions on a patient manifold (Figure 5B). There, we see that cluster proportions arranged on this manifold reveal clusters that are changing across the space. This analysis indicates clearly that cluster 1 is representative of acute subjects and cluster 5 is representative of the healthy subjects. Furthermore, we can evaluate the same individual when measured after acute infection, and then later at a convalescent time point (Figure 5C). Viewed in this way, we see that cluster 11 is also more present in most subjects when they came in with an acute infection than at the convalescent time point.

**Figure 5:**
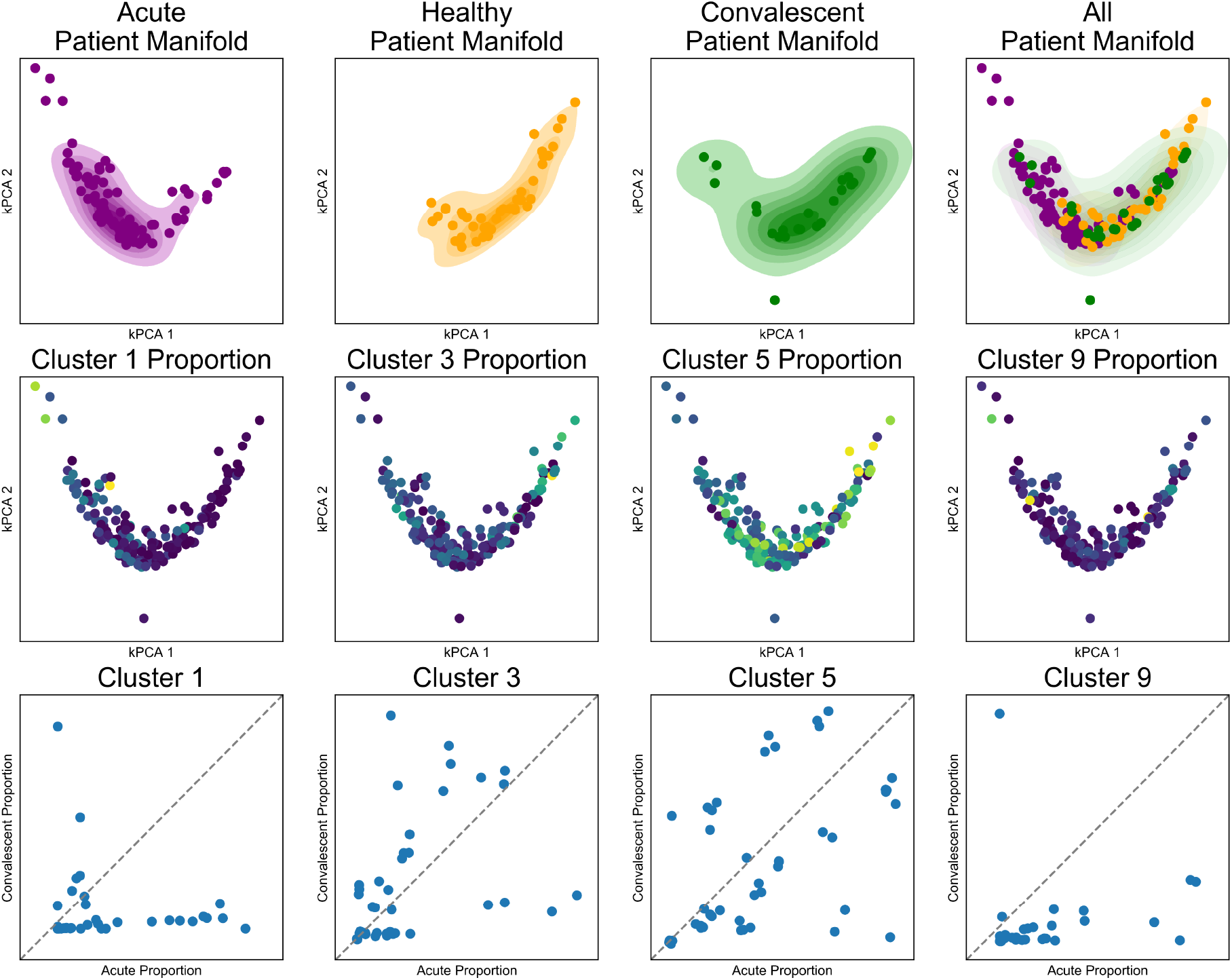
SAUCIE produces patient manifolds from single-cell cluster signatures. Top row) The patient manifold identified by SAUCIE cluster proportions, visualized by kernel PCA with acute, healthy, convalescent, and all subjects combined from left to right. The healthy manifold overlaps with the convalescent manifold to a much higher degree than the acute manifold. Middle row) The same patient manifold shown colored by each patient’s cluster proportion. Cluster 1 is more prevalent in acute, cluster 3 in healthy, cluster 5 is ubiquitous, and cluster 9 is rare and in acute patients. Bottom row) A comparison of the cluster proportion for acute (X-axis) versus convalescent (Y-axis) for patients that have matched samples.

#### 2.3.3 Visualization

SAUCIE can process all cells from all subjects to construct a cellular manifold and extract its features. First, we visualize this manifold using the 2-D visualization layer. Figure 6A is divided into two embeddings that show the cell manifolds for acute and healthy subjects separately. As can be seen, there is a characteristic change in the manifold that becomes apparent when comparing the embeddings side-by-side. The acute subjects have cell populations distinctly missing that are present in the healthy subjects.

**Figure 6:**
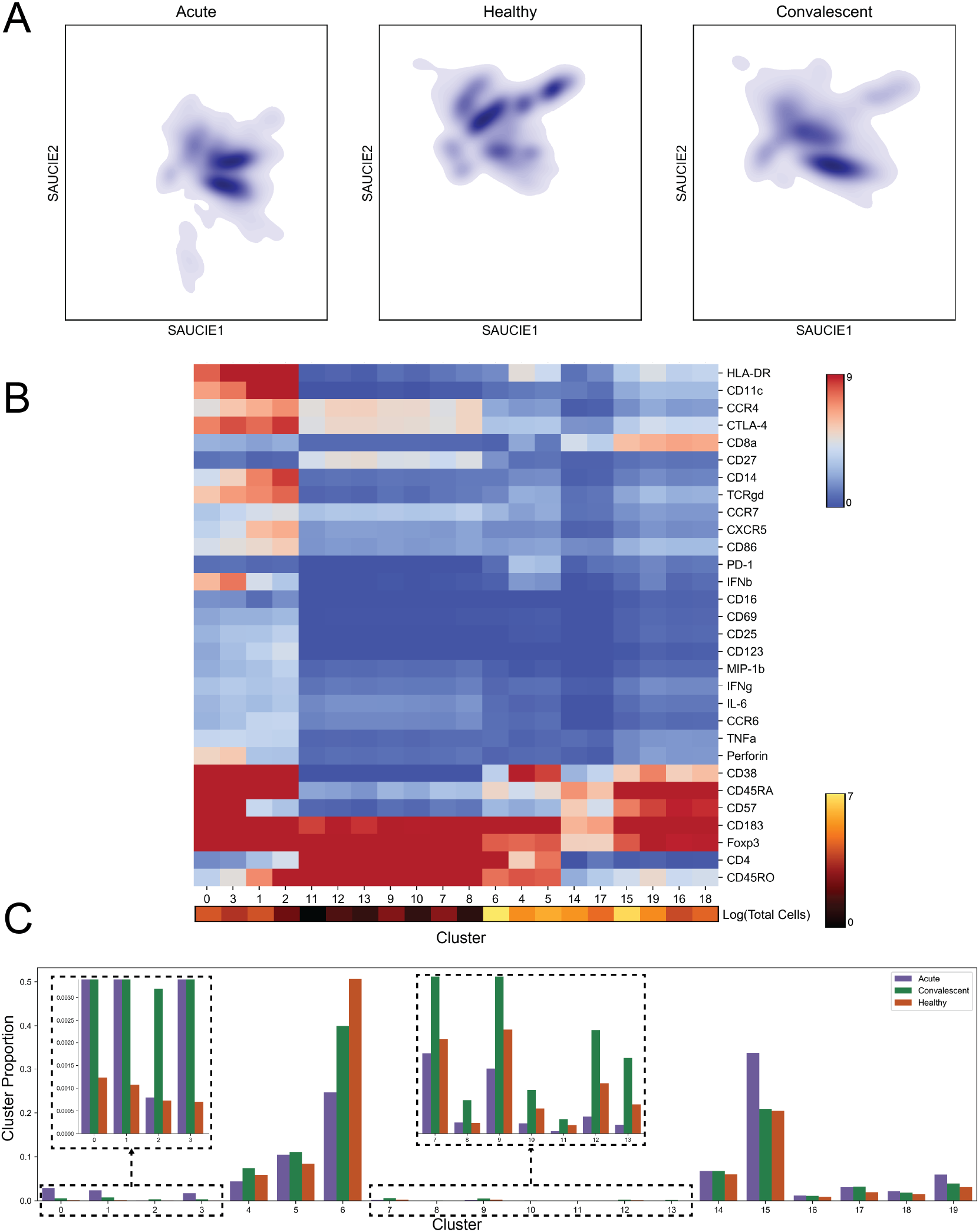
SAUCIE identifies and characterizes cellular clusters, whose proportions can be used to compare patients. A) The cell manifolds identified by the two-dimensional SAUCIE embedding layer for the T lymphocyte subsets from acute, healthy, and convalescent subjects. B) A heatmap showing clusters along the horizontal axis and markers along the vertical axis. Cluster sizes are represented as a color bar beneath the heatmap. C) Cluster proportions for acute, convalescent, and healthy patients.

After characterizing the nature of the cellular space in the aggregate, we can additionally analyze manifolds formed by the distributions of T lymphocytes within each patient separately. As each patient has a heterogeneous population of cells, including with different total numbers of cells, it becomes a challenge to define a meaningful measure of similarity between the individuals. Here we are able to leverage the manifold constructed by the SAUCIE embedding and calculate MMD (a distribution distance) between the distribution of cells in the latent space for each pair of subjects. With a measure of similarity between each pair of patients, we can now construct a manifold not of the cells but also of the *subjects* (Figure 5A).

#### 2.3.4 Comparison to existing method

We next compare the SAUCIE pipeline of batch correcting, clustering, and visualizing single-cell data from a cohort of subjects to an alternative approach called metaclustering [47]. We first cluster each sample individually with Phenograph. Then, we represent each cluster as its centroid and use Phenograph again on the clusters to obtain metaclusters. We examine the pipelines on ten of the 180 samples here, where the metaclustering approach took forty minutes. We note that the SAUCIE pipeline took *45 minutes* to process all 180 samples, while the metaclustering approach would take *12 hours* to process all of them. Figure S10 shows tSNE embeddings of the cluster centroids where the size of the cluster is proportional to the size of the point. Coloring by sample, we see that the metaclusters have identified batch effects. Metacluster 0 is only composed of samples 1, 3, 4, and 5. These samples have no clusters in any other metacluster, and none of the other samples have any cluster in this metacluster. Examining the gene expression heatmap, we see that metacluster 0 has separated cells with high CD86 values, which were shown earlier to be batch effects. Moreover, the metaclusters are very heterogeneous internally with respect to gene expression. This is a results of metaclustering the cluster centroids, as the metaclusters then have no information about the individual cells comprising that centroid.

In contrast, Figure S11 shows the SAUCIE pipeline on these ten samples. The cluster proportions show that each cluster is fully mixed with respect to the samples, as opposed to the sample-segregated metaclusters of the previous approach. Similarly, the clusters are more homogeneous internally, meaning they actually keep similar cells together, as opposed to the metaclusters, which lost this information when each cluster was represented by only its centroid. Finally, we find that SAUCIE effectively compares cells across subjects, while the metaclustering approach still fails at patient-to-patient comparisons, instead only identifying batch effect variation. This emphasizes the importance of multitask learning using a unified representation in SAUCIE.

## 3 Discussion

We presented SAUCIE, a neural network framework that streamlines exploratory analysis of datasets that contain a multitude of samples and a large volume of single cells measured in each sample. The key advantage in SAUCIE is its ability to perform a variety of crucial tasks on single-cell datasets in a highly scalable fashion (utilizing the parallelizability of deep learning with GPUs) without needing to call external algorithms or processing methods. As a result, SAUCIE is able to process multisample data in a unified way using a single underlying representation learned by a deep autoencoder. Thus, different samples can be visualized in the same coordinates without batch effects via the embedding layer of the neural network, and cluster proportions can be directly compared, since the whole dataset is decomposed into a single set of clusters without requiring cluster matching or metaclustering. These unified representations can be readily used for inter-sample comparisons and stratification, on the basis of their underlying cell-to-cell heterogeneity.

Mathematically, SAUCIE presents a new way of utilizing deep learning in the analysis of biological and biomedical data by directly reading and interpreting hidden layers that are regularized in novel ways to understand and correct different aspects of data. Thus far, deep learning has primarily been used in biology and medicine as a black-box model designed to train classifiers that often mimic human classifications of disease or pathology. However, the network internal layers themselves are typically not examined for mechanistic understanding. SAUCIE is leading a new wave of deep learning models that obtain information from internal layers of a deep network. Deep autoencoding neural networks essentially perform nonlinear dimensionality reduction on the data. As such they could be used “off-the-shelf” for obtaining new coordinates for data in a reduced-dimension space, to which other algorithms can be applied. However, in SAUCIE we aim to go further to structure the reduced dimensions in specifically interpretable ways using novel regularizations. Our information-theoretic regularization encourages near-binary activations of an internal layer, thus making the layer amenable to directly output encoded cluster identifications. We believe that this is just the first foray into what could be a vast number of such regularizations that can offer interpretability of specialized layers in neural networks, thus turning these “black boxes” into “glass boxes.”

The ability to stratify patients on the basis of their single-cell subpopulations, which can emerge as features in deep neural networks, can be key to a new generations of biomarkers that can be used in diagnosis and treatment. Traditionally, biomarkers are proteins or antibodies that are circulating in blood, which signals the presence of infection or other conditions. However, immune cells are highly plastic and can evolve or activate in specific ways in response to disease conditions in different patients. Here, we showcase the heterogeneity of immune cells in response to acute dengue infection in a large patient cohort. We see that specific subpopulations are enriched in the acute conditions, as opposed to convalescent or healthy controls. We showed with our dengue dataset it is possible discover cell populations, even rare ones that are indicative of patient and experimental conditions. Other datasets comprising of large patient cohorts measured at single-cell resolution are underway already in many hospitals and clinical trials. In the future, we are confident that this capability will be useful in many studies including immunotherapy, autoimmunity, and cancer, where there are immune subsets that emerge in response.

## 4 Methods

### 4.1 Computational Methods

In this section we explain the SAUCIE framework in greater detail including the philosophy behind using autoencoders for learning the cellular manifold, details of the regularizations used in different layers of SAUCIE to achieve particular data analysis tasks as well as training and implementation details. Finally, we discuss the emergent higher level organization of the patient manifold as a result of the cellular manifold of the subjects learned by SAUCIE.

#### 4.1.1 Multitask manifold learning

A popular and effective approach for processing big high-dimensional data in genomics, as well as other fields, is to intuitively model the intrinsic geometry of the data as being sampled from a low dimensional manifold – this is commonly referred to as the manifold assumption [17]. This assumption essentially means that local regions in the data can be linearly mapped to low dimensional coordinates, while the nonlinearity and high dimensionality in the data comes from the curvature of the manifold. Typically, a notion of locality is derived from the data with nearest-neighbor search or adaptive kernels to define local neighborhoods that can approximate tangent spaces of the manifold. Then, these neighborhoods are either used directly for optimizing low dimensional embeddings (e.g., in TSNE [48] and LLE [49]), or they are used to infer a global data manifold by considering relations between them (e.g., using diffusion geometry [14,15,50,51]). In the latter case, the data manifold enables several applications, including dimensionality reduction [14,51], clustering [50,52–54], imputation [15], and extracting latent data features [55–57].

The characterization of the intrinsic data geometry as a data manifold is also closely related to the underlying approach in SAUCIE. Indeed, neural networks can be considered as piecewise linear approximations of target functions [58]. In our case, we essentially approximate the data manifold coordinate charts and their inverse with the autoencoder architecture of SAUCIE. The encoder training identifies local patches and maps them to low dimensional coordinates, while sewing these patches together in this embedding to provide a unified visualization. The decoder learns the linear relation between these intrinsic coordinates and the tangent spaces of the manifold, positioned in the high dimension. This also results in a projection of data points on the manifold (via its tangent spaces), which creates a denoising effect similar to the diffusion-based one used recently in MAGIC [15]. Finally, the clustering layer in SAUCIE is trained to recognize and aggregate similar data regions to ensure an appropriate granularity (or resolution) of the identified neighborhoods and prevent excessive fragmentation of the manifold. For more discussion regarding the relations between deep learning and manifold learning we refer the reader to [2,16,59].

While tools using the scaffold of manifold learning have emerged for various tasks in single cell data analysis, there is currently no unified manifold model that provides all of the necessary tasks in a scalable fashion. For example, MAGIC [15] uses manifold learning to impute the data, but does not address embedding, visualization, or clustering. Diffusion pseudotime [55] provides an organization of the data to infer latent temporal structure and identifies trajectories, but it does not deal with imputation, clustering, or visualization. Furthermore, manifold learning methods do not work well across batches and typically just focus on single batches. Thus, their construction may suffer from batch effects and be dominated by the geometry between batches rather than their biology, as demonstrated by the example of Phenograph in Figure S10.

To address these shortcomings, SAUCIE performs all operations on a unified manifold geometry, which is learned implicitly by a deep multitasking neural network. It utilizes the scalability of deep learning to process high throughput data and construct a manifold that is jointly optimized for multiple tasks; namely, clustering, visualization, imputation, and batch correction. Therefore, the tasks themselves respect the manifold assumption and have the associated advantages, such as robustness to noise, while also agreeing with each other on a coherent underlying structure of the data.

#### 4.1.2 SAUCIE architecture

SAUCIE consists of three encoding layers, an embedding layer, and then three decoding layers. The default number of neurons per hidden layer in the encoder used were 512, 256, and 128 with a symmetric decoder. The GMM dataset, being simpler, was clustered with layers of 50, 30, and 10. For batch correction, the best results were achieved with layer sizes of 1024, 512, and 256. The ID regularization was applied to the final decoder layer, which uses a ReLU. The two-dimensional embedding layer uses a linear activation, while all other layers use a leaky rectified linear activation with 0.2 leak. The coefficients *λ*_*d*_ and *λ*_*c*_ were chosen depending on the dataset, with the best values generally being *λ*_*d*_ twice *λ*_*c*_. Their magnitude was guided by the effect of these two knobs on the granularity (shown in Figure S9). Training was performed with minibatches of 256, mean-squared-error for the reconstruction error function, and the optimizer chosen is ADAM with learning rate 0.001.

#### 4.1.3 Batch correction and MMD Regularization

A major challenge in the analysis of single-cell data is dealing with so-called batch effects that result from technical variability between replicates of an experiment. Combining replicates often results in technical and experimental artifacts being the dominant source of variability in the data, even though this variability is entirely artificial. This experimental noise can come in the form of dropout, changes of scale, changes of location, or even more complicated differences in the distributions of each batch. It is infeasible to parametrically address all of the potential differences explicitly, for example, by assuming measurements are drawn from a Gaussian distribution. Instead of addressing specific explicit models of noise, SAUCIE minimizes a distance metric between distributions. The batch correction term *L*_*b*_ calculates the Maximal Mean Discrepancy (MMD) [60] between batches, as

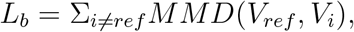

where *V_ref_* is the visualization layer of one of the replicates, arbitrarily chosen to be considered as a reference batch. MMD compares the average distance from each point to any other point in its own batch, with the distance to points in the other batch. MMD is zero only when two distributions are equal. Thus minimizing this metric encourages SAUCIE to align the batches. MMD has been used effectively to remedy batch effects in residual networks, but here SAUCIE uses it in a feedforward autoencoder and combines it with other tasks of interest in biological exploratory data analysis [21].

The choice of reference does not affect the degree to which two distributions can be aligned, but a reference batch is necessary because the encoding layers of a standard network will be encouraged to embed different batches in different places in the visualization layer. It does this because the decoder is required to make its reconstruction 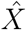 match the original data in *X*, which includes the batch effects. To remedy this, the decoder in SAUCIE is required to reconstruct the reference batch exactly as usual, but other batches must only be reconstructed to preserve the points normalized by mean and variance. Consequently, the MMD regularization term will be minimized when batches are aligned, and the decoder need only be able to reconstruct the exact values of the reference batch and the *relative values* of the non-reference batches. The non-reference batches will be aligned to the reference batch in a way that preserves their internal structure as best as possible.

#### 4.1.4 Regularizations and Post-processing for clustering

##### Information Dimension Regularization

We consider the task of clustering data points by interpreting the sparse layer *B* in the network as encoding cluster assignments. We note that a common activation function used to introduce nonlinearities in neural networks (including SAUCIE) is the Rectified Linear Unit (ReLU), and it provides a natural threshold for binarizing neuron activation to be either zero or one. These units are either “off” at or below zero or “on” for any positive value, so a small positive value *ϵ* can be used a threshold to binarize the activations in *B*. This results in an interpretable clustering layer that creates ‘digital’ cluster codes out of an ‘analog’ hidden layer, thus providing a binary code for each input point of the network. These binary codes are in turn used as cluster identifiers in order to group data points with the same code into a single cluster.

In order to automatically learn an appropriate granularity of clusters, we developed a novel regularization that encourages near-binary activations and minimizes the information (i.e., number of clusters) in the clustering layer. Our regularization is inspired by the von Neumann (or spectral) entropy of a linear operator [61], which is computed as the Shannon entropy of their normalized eigenvalues [62,63]. This entropy serves as a proxy for the numerical rank of the operator [51], and thus provides an estimation of the essential dimensionality of its range. In our case, we extend this notion to the nonlinear transformation of the neural network by treating neurons as our equivalent of eigenvalues, and computing the entropy of their total activation over a batch. We call this entropy ‘information dimension’ (ID) and the corresponding ID regularization aims to minimize this entropy while still encoding sufficient information to allow reconstruction of the input data points.

The ID regularization is computed from the clustering layer activations in *B* by first computing the activation of each neuron *j* as 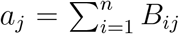, then normalizing these activations to form an activation distribution 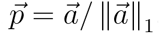, and finally computing the entropy of this activation distribution as

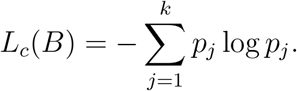

By penalizing the entropy of neuron activations, this regularization encourages a sparse and binary encoding. This counters the natural tendency of neural networks to maximize the amount of captured (i.e., encoded) information by spreading activations out across a layer evenly. By forcing the activations to be concentrated in just a few distinct neurons, different inputs end up being represented with rather similar activation patterns, and thus naturally clustered. When combined with the reconstruction loss, the network will retain enough information in the sparse layer for the decoder to reconstruct the input, keeping similar points in the same cluster.

##### Intracluster distance regularization

The digital codes learned by SAUCIE create an opportunity to interpret them as clusters, but these clusters would not necessarily be comprised of only similar points. To emphasize that inputs only be represented by the same digital code if they are similar to each other, SAUCIE also penalizes intracluster pairwise distances. Beyond suffering reconstruction loss, using the same code for points that are far away from each other will now incur an even greater loss.

This loss is calculated as the euclidean distance between points with the same binary code:

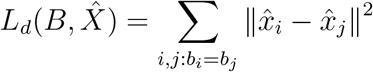

where 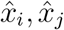 and *b*_*i*_, *b_j_* are the *i*-th and *j*-th rows of 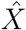 and *B*, respectively.

Since ID regularization is minimized by using the same code to represent all inputs, this term acts as an opposing balance. Intracluster distances are minimized when all points are in a cluster by themselves. Together with the reconstruction penalty, these terms encourage SAUCIE to learn clusters that are composed of as many points as possible that are near to each other.

An additional benefit of clustering via regularization is that not only is the number of clusters not needed to be set *a priori*, but by changing the value of *λ*_*c*_ the level of granularity of the clustering can be controlled, so both coarse clustering and fine clustering can be obtained to further add insight into the underlying structure of the data.

##### Cluster merging

As the binarized neural network may not converge to the ideal level of granularity due to the many possible local optima in the loss landscape, we process the SAUCIE clustering with a cluster merge step to fix the ideal level of granularity everywhere. The cluster merging is performed by calculating MMD between clusters in the SAUCIE latent space and merging all clusters *i, j* ∈ *C*, where *C* is the set of all clusters, such that both of the following equations hold

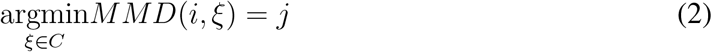

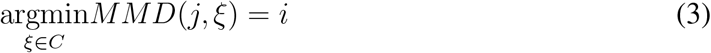

This merging finds clusters that would be a single cluster in another granularity and fixes them to a single cluster.

#### 4.1.5 Patient Manifold Visualization

In addition to the cell-level manifold constructed by SAUCIE, we also consider the geometry between samples to provide a coarser patient-level manifold. We construct and embed this manifold in low dimensions by applying kernel-PCA (kPCA) [64] with an RBF kernel to the metric space defined by MMD distances between subjects. This augments the analysis SAUCIE provides of the biological variations identified in the cell space with an analysis of the variation in the patient space. Normally, without batch correction, the two sources of variation would be confounded, and batch effects would prevent clear analysis at either level (patient or cell) across batches. With our approach here we are able to separate them to provide on one hand, a stable (batch-invariant) cell-level geometry by the SAUCIE embedding, and on the other hand, a robust patient geometry provided by kPCA embedding. The patient geometry then allows us to recover patient-level differences and utilize them further for data exploration, in conjunction with the cell-level information. For example, as Figure 5A shows, we have a notable stratification between the acute and non-acute subjects. There is also a noticeable difference between the convalescent subjects and the acute, albeit a less drastic one than the difference between acute subjects and the others.

#### 4.1.6 Training

To perform multiple tasks, SAUCIE uses a single architecture as described above, but is run and optimized sequentially. The first run imputes noisy values and corrects batch effects in the original data. This preprocessed data is then run through SAUCIE again to obtain a visualization and to pick out clusters. The different runs are done by optimizing different objective functions. In the following, we describe the optimization of each run over a single batch of *n* data points. However, the full optimization of each run independently utilizes multiple (mini-)batches in order to converge and minimize the described loss functions.

For the first run, formally let *X* be an *n* × *d* input batch, where each row is a single data point, and *d* is the number of features in the data. It is passed through a cascade of encoding linear and nonlinear transformations. Then, a cascade of decoding transformations reconstruct the denoised batch 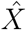, which has the same dimensions as the input *X* and is optimized to reconstruct it.

For the next run, the cleaned batch 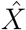 is passed through encoding transformations and a visualization layer denoted by *V* ∈ ℝ^*n*×2^. We also consider a clustering layer in another run where the decoder outputs near-binary activations *B* ∈ ℝ^*n*×*d_B_*^, where *d*_*B*_ is the number of hidden nodes in the layer, which will be used to encode cluster assignments, as described below. The activations in *B* are then passed to the reconstruction 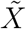 that has the same dimensions as 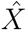 (and *X*) and is optimized to reconstruct the cleaned batch.

The loss function of all runs starts with a reconstruction loss *L_r_* forcing the autoencoder to learn to reconstruct its input at the end. SAUCIE uses the standard mean-squared error loss (i.e., 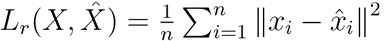, where *x*_*i*_ and 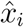 are the *i*-th row of *X* and 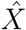 correspondingly). We note that while MSE is a standard and effective choice in general, other loss functions can also be used here as application-specific substitutes that may be more appropriate for particular types of data. For the first run, we add to this loss a regularization term *L_b_* that enables SAUCIE to perform batch correction. This regularization is computed from the visualization layer to ensure consistency across subsampled batches. The resulting total loss is then

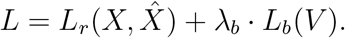

The loss function of the clustering run then optimizes *L*_*r*_ along with two regularization terms *L*_*c*_ and *L_d_* that together enable SAUCIE to learn clusters:

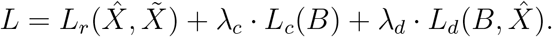

The first term *L*_*c*_ guides SAUCIE to learn binary representations via the activations in *B* using a novel information dimensionality penalty that we introduce in this paper. The second term *L*_*d*_ encourages interpretable clusters that contain similar points by penalizing intra-cluster distances in the cleaned batch 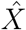, which is fixed for this run.

#### 4.1.7 Runtime Comparison Methodology

For each visualization, clustering, and imputation method, the dataset of size *N* was given to the method as input and returned the appropriate output. For batch correction, the dataset of size *N* was divided into two equal-sized batches that were corrected. For the methods that operated on minibatches, minibatches of size 128 were used. For the methods that train by stochastic gradient descent, the number of steps was determined by taking the total number of points and dividing by the size of the minibatch, so that a complete pass through the entire dataset was performed. In order to return clusters, the latent space of scVI must be clustered by another method, and since the number of clusters is not known ahead of time, the fastest method that does not require this to be known (Phenograph) was used. For SAUCIE, batch correction, imputation, clustering, and visualization were all produced in the timed run. All computations were performed on a single machine with 16 CPU cores and a GeForce GTX 1080 GPU.

#### 4.1.8 Number of Clusters

As discussed earlier, the number of clusters resulting from SAUCIE is not specified in advance, but dictated by the structure of the data that the model discovers, and by the choice of regularization coefficients *λ*_*d*_ and *λ*_*c*_. For a given value of *λ*_*d*_, as *λ*_*c*_ increases, the number of clusters decreases. Increasing *λ*_*d*_, on the other hand, increases the number of clusters (Figure S9). This is because *λ*_*c*_ penalizes entropy in the activations of the *n* neurons in the clustering layer of the network. While entropy can be initially decreased by making all *n* neurons either 0 or 1, it can be further decreased by making all *n* neurons 0. Thus, as this term is considered more influential in the total loss, in the extreme, all points can be mapped to the same binary code. In contrast, *λ*_*d*_ penalizing intra-cluster distances, so this value can be decreased by making clusters smaller and smaller (and thus getting more of them). In the extreme for this term, every point can be made its own cluster and intra-cluster distances would decrease to 0. By balancing these two, the desired granularity of clustering can be obtained from SAUCIE. In our experiments, we find making *λ*_*d*_ to be between two and three times larger than *λ*_*c*_, with values around 0.2 generally results in medium coarse-grained clustering. Another consideration that affects the number of clusters is the number of neurons in the clustering layer. We found varying this number does not improve performance and for all experiments here we use a fixed size of 256 neurons.

### 4.2 Experimental methods

#### 4.2.1 Study Subjects

Dengue patients and healthy volunteers were enrolled with with written informed consent under the guidelines of the Human Investigations Committees of the NIMHANS and Apollo Hospital, and Yale University [19]. The Human Investigations Committee of each institution approved this study. Patients with dengue virus infection were defined as dengue fever using WHO-defined clinical criteria, and/or laboratory testing of viral load or serotyping at the time of infection. Healthy volunteers included household contacts of dengue patients present in the same endemic area. Participants were of both genders (26.7% female) and were all of Indian heritage. Subjects from the symptomatic and healthy groups were not statistically different for age, gender, or race in this study.

#### 4.2.2 Sample Collection and Cell Isolation

Heparinized blood was collected from patients and healthy volunteers and employed a 42 marker panel of metal conjugated antibodies following methods previously described [65,66]. Purification of peripheral blood mononuclear cells (PBMCs) was performed by density-gradient centrifugation using Ficoll-Paque (GE Healthcare) according to the manufacturer’s instructions following isolation and cryopreservation guidelines established by the Human Immunology Phenotyping Consortium. PBMCs for CyTOF were frozen in 90% FBS containing 10% DMSO and stored in liquid N2 for shipping following the guidelines of the DBT. Samples for this study were received in three shipments and viability was average 85% (range 50–98) across the dates.

#### 4.2.3 Mass Cytometry Acquisition

For mass cytometry at Yale University, PBMCs (5 × 106 cells/vial) were thawed incubated in Benzonase (50U/ml) in RPMI/10% human serum, and seeded in 96-well culture plate (6 × 103-1.2 × 106 cells/well. Monensin (2μΜ, eBioscience) and Brefeldin A (3μg/ml, eBioScience) added for the final 4 h of incubation for all groups. Groups of samples (8-13/day) were infected in vitro per day on 5 separate days and included a CD45-labeled spike-in reference sample in every sample. Surface markers were labeled prior to fixation and detailed staining protocols have been described. Briefly, cells were transferred to 96-well deep well plates (Sigma), resuspended in 25 μΜ cisplatin (Enzo Life Sciences) for one minute, and quenched with 100% FBS. Cells were surface labeled for 30 min on ice, fixed (BD FACS Lyse), and frozen at –80°C. Intracellular labeling was conducted on batches of cells (12/day). Fixed PBMCs were permeabilized (BD FACS Perm II) for labeling with intracellular antibodies for 45 min on ice. Cells were suspended overnight in iridium interchelator (125 nM; Fluidigm) in 2% paraformaldehyde in PBS and washed 1X in PBS and 2X in H2O immediately before acquisition. A single batch of metal-conjugated antibodies was used throughout for labeling panels. Metal-conjugated antibodies were purchased from Fluidigm, Longwood CyTOF Resource Core (Cambridge, MA), or carrier-free antibodies were conjugated in house using MaxPar X8 labeling kits according to manufacturer’s instructions (Fluidigm). A total of 180 samples were assessed by the Helios (Fluidigm) on 15 independent experiment dates using a flow rate of 0.03 ml/min in the presence of EQ Calibration beads (Fluidigm) for normalization. An average of 112, 537 ± 71, 444 cells (mean ± s.d.) from each sample were acquired and analyzed by CyTOF. Data was preprocessed with the hyperbolic sine transformation. Additional experimental details will be given in [19].

### 4.3 Grant Support

This work was supported in part by awards from the NIH (AI089992), the Indo-U.S. Vaccine Action Program. It was also supported by the CZI grant for computational tools.

## 5 Software

SAUCIE is written in Python using the Tensorflow library for deep learning. The source code is available at https://github.com/KrishnaswamyLab/SAUCIE/.

---

**Supplemental Figure S1:**
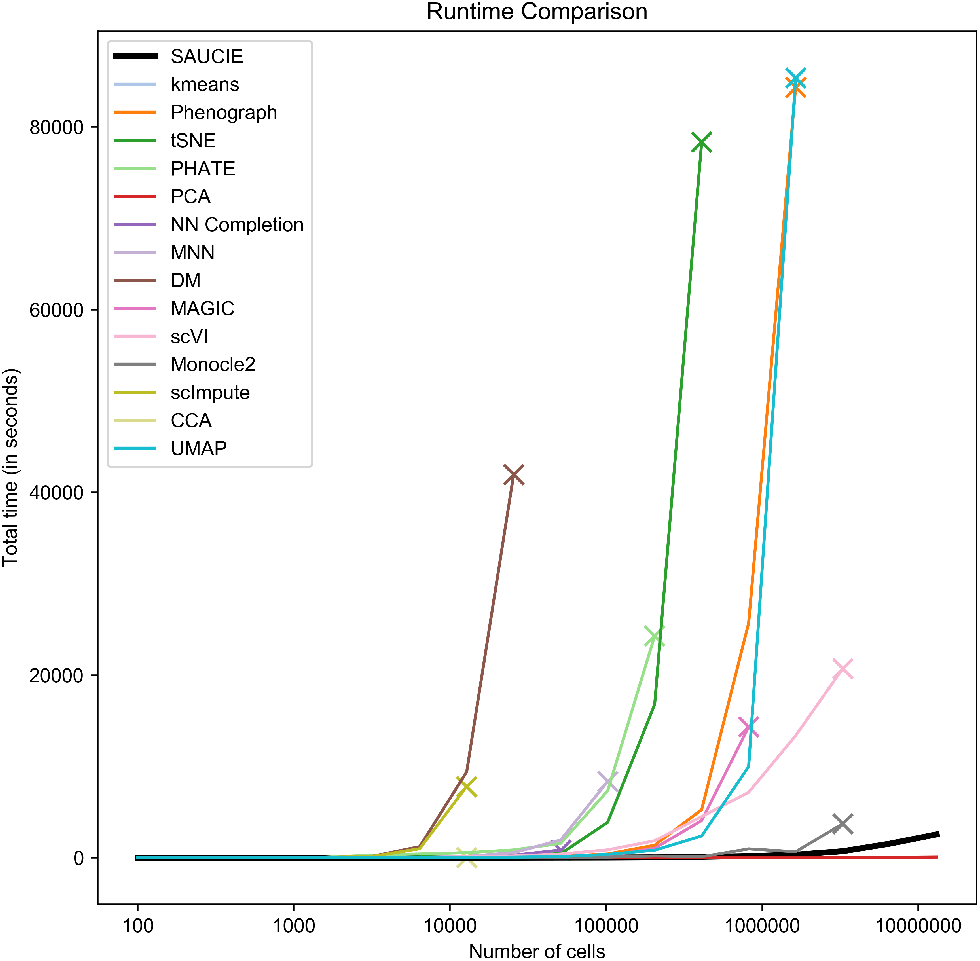
Comparison of runtimes on an increasing number of points. The number of points is represented on the horizontal axis and the time in seconds the method took to complete is on the vertical axis. If a method ran out of resources and could not complete a run for a certain number of points, that is demarcated with an ‘x’ and no further time points were attempted for that method. SAUCIE is the fastest method besides PCA and kmeans.

**Supplemental Figure S2:**
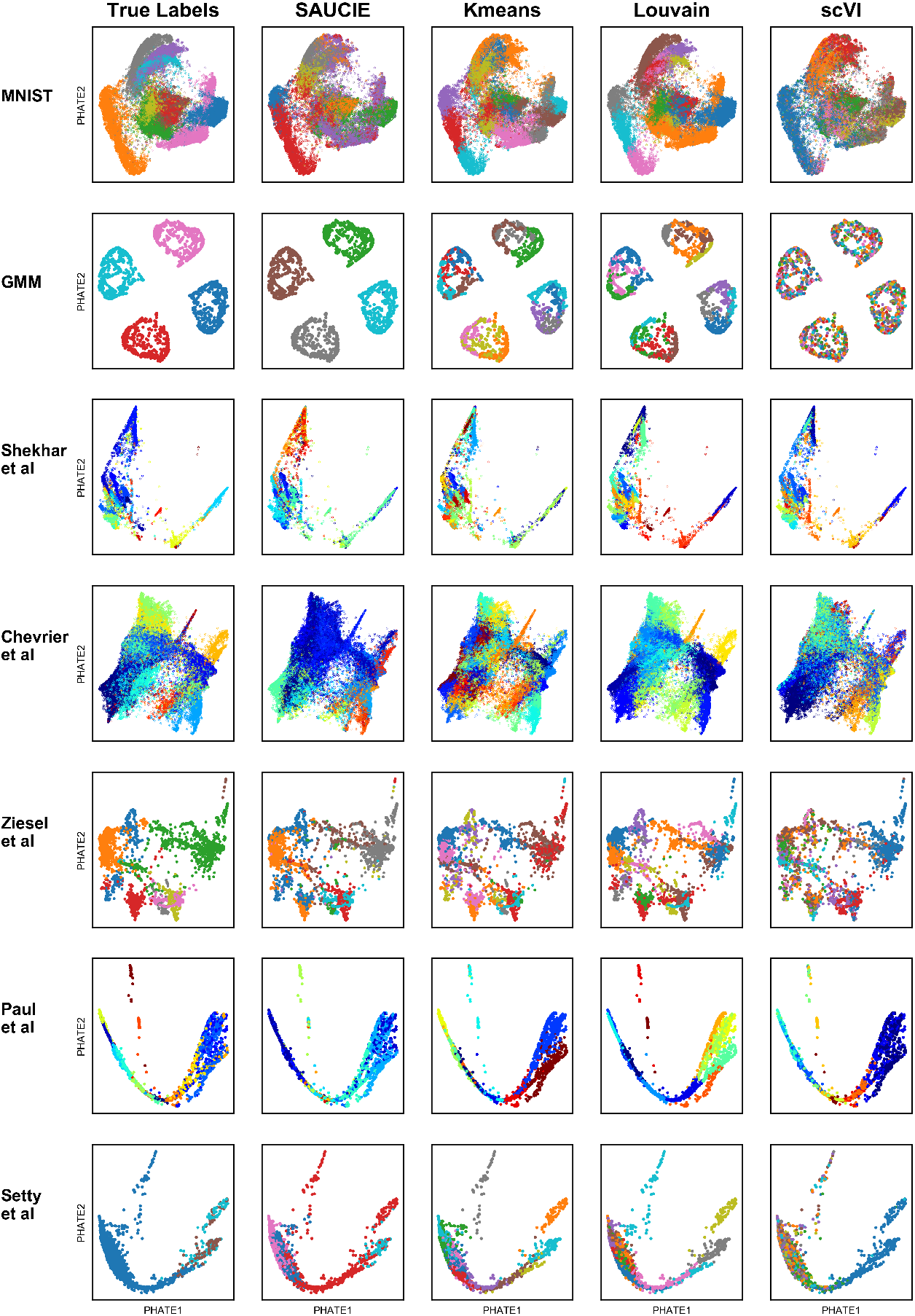
A comparison of the SAUCIE clustering to other clustering methods on artificial and real data. Rows show the different datasets. Along with the first two artificial datasets, there are two CyTOF datasets and three scRNA-seq datasets. Columns show the different clustering methods. From left to right: True “ground truth” labels, SAUCIE, kmeans, Phenograph, scVI.

**Supplemental Figure S3:**
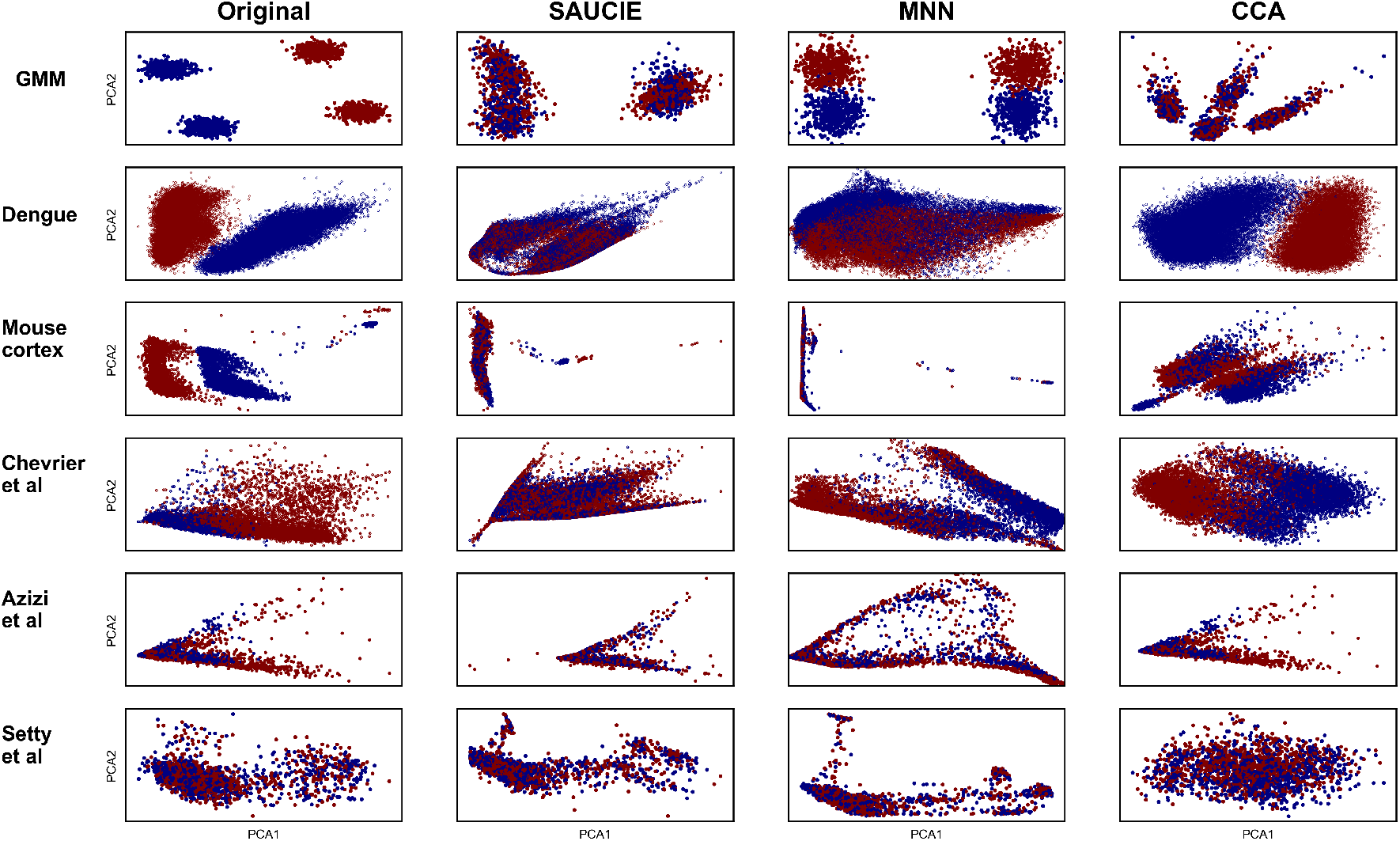
A comparison of batch correction with SAUCIE to other methods on an artificial dataset, two technical replicates from the dengue CyTOF data, non-technical replicates on scRNA-seq batches from mouse cortex, and then public data from Chevrier et al, Azizi et al, and Setty et al. Rows show the different datasets. Columns show the different batch correction methods. From left to right: The original data prior to batch correction, SAUCIE, mutual nearest neighbors (MNN), canonical correlation analysis (CCA).

**Supplemental Figure S4:**
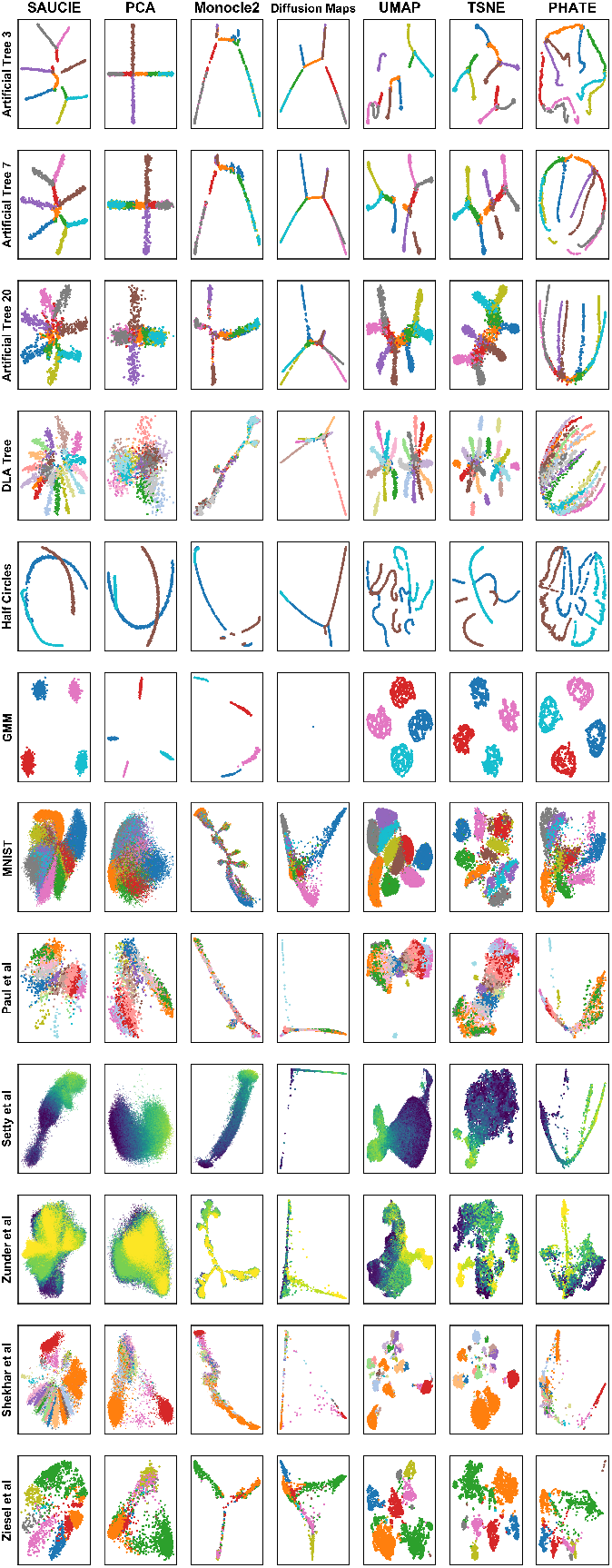
A comparison of the SAUCIE visualization to other methods on a number of artificial and real datasets. The columns show the different methods. From left to right: SAUCIE, PCA, Monocle2, Diffusion Maps, UMAP, tSNE, PHATE. The rows show the different datasets. From top to bottom: Artificially generated trees with varying amounts of noise, random tree generated with diffusion limited aggregation (DLA), intersecting half circles, Gaussian mixture model, MNIST, scRNA-seq hematopoiesis from Paul et al. 2015 [27], CyTOF T cell development from Setty et al. 2016 [22], CyTOF ipsc from Zunder at al. 2016 [25], scRNA-seq retinal bipolar cells from Shekhar et al. 2016 [26], scRNA-seq mouse cortex from Zeisel et al. 2015 [28].

**Supplemental Figure S5:**
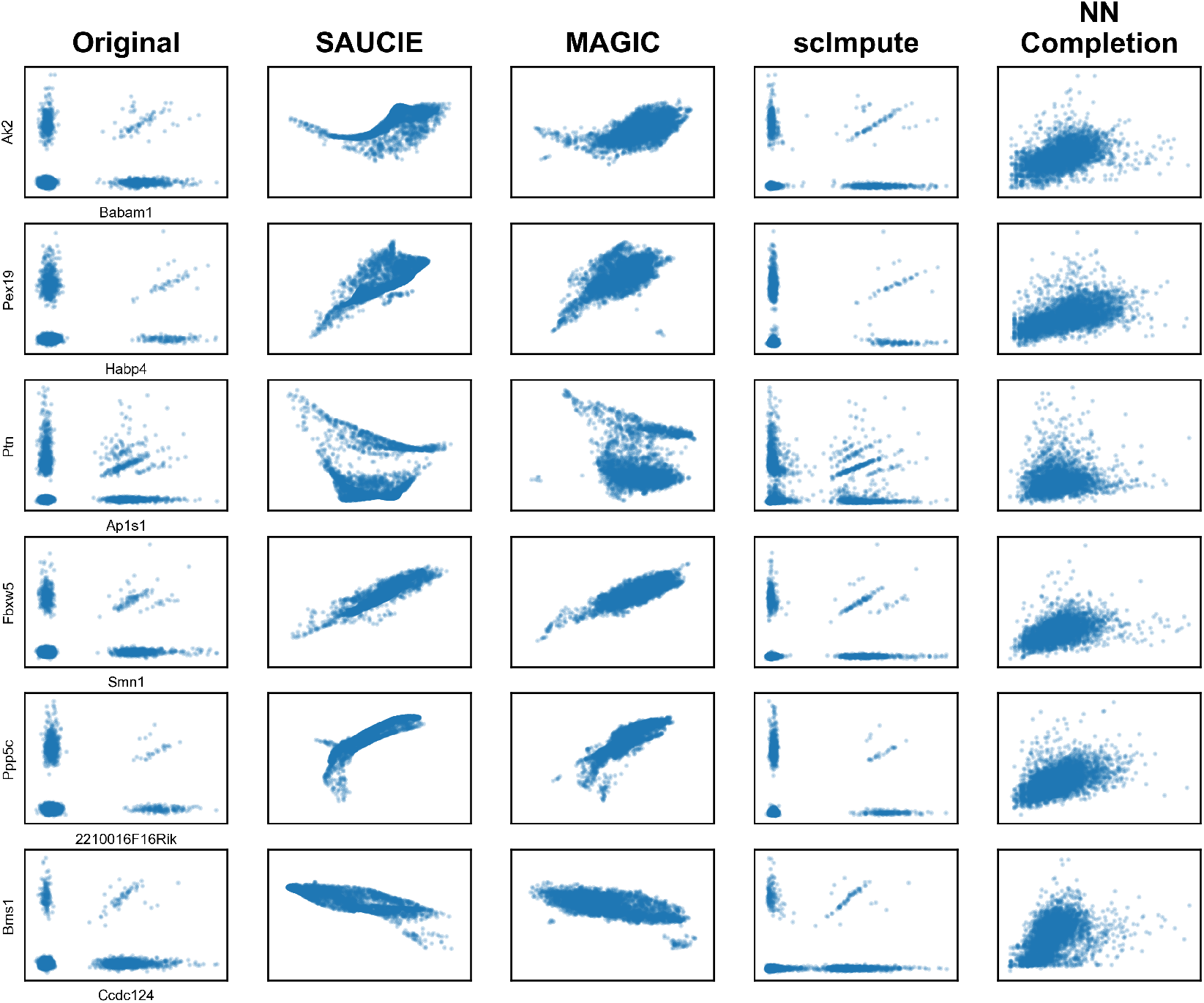
A comparison of imputation methods including SAUCIE. Several gene-gene associations are shown from the 10x mouse cortex dataset. From left to right: The original (sparse) data, data after imputation with SAUCIE, MAGIC, scImpute, and nearest neighbor completion.

**Supplemental Figure S6:**
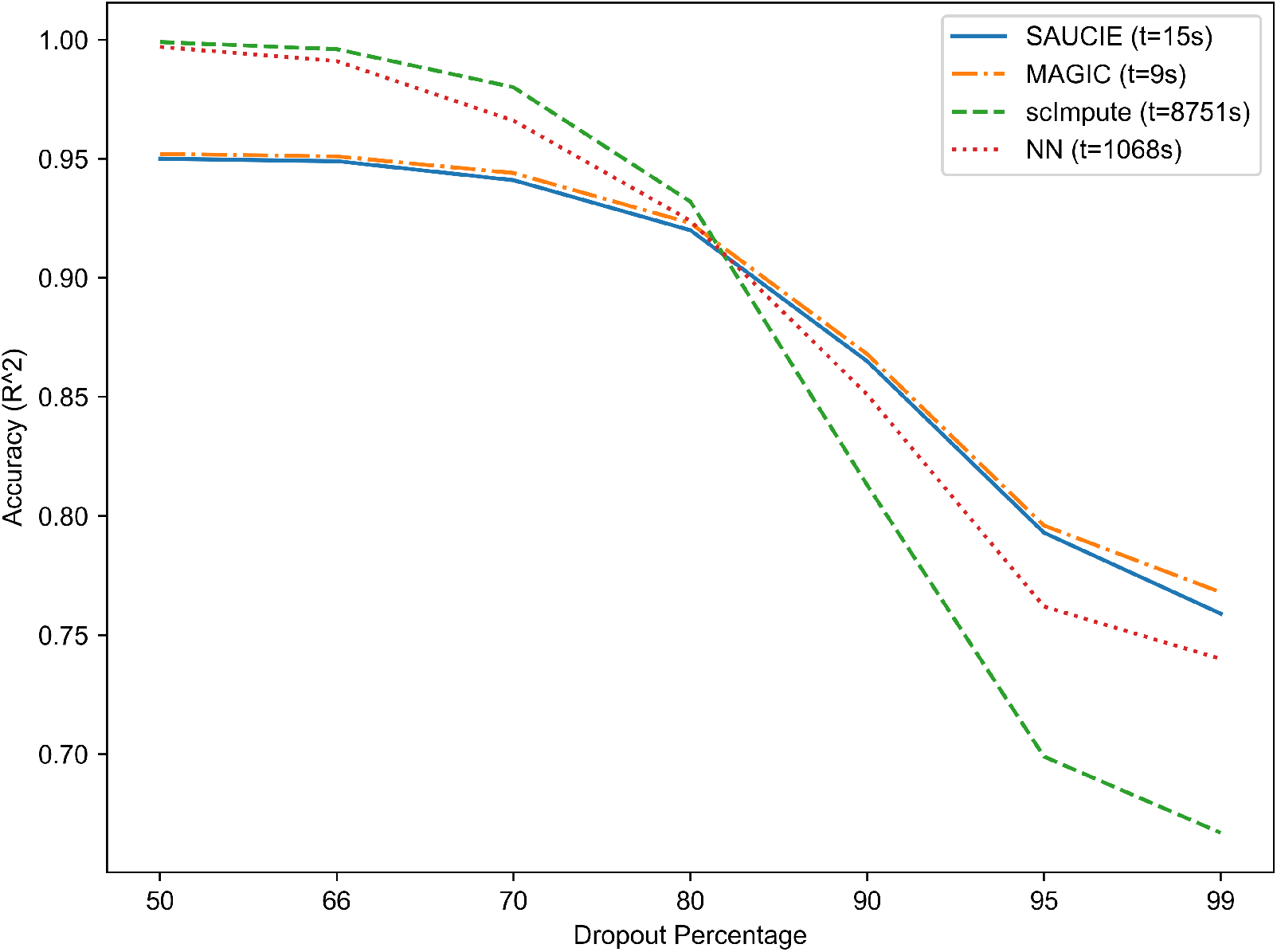
A comparison of imputation with SAUCIE to other methods on the simulated dropout experiment. Increasing amounts of dropout are along the horizontal axis from left to right, and the accuracy of each method as measured by *R*^2^ is along the vertical axis. The time each method took to complete is in the legend in seconds.

**Supplemental Figure S7:**
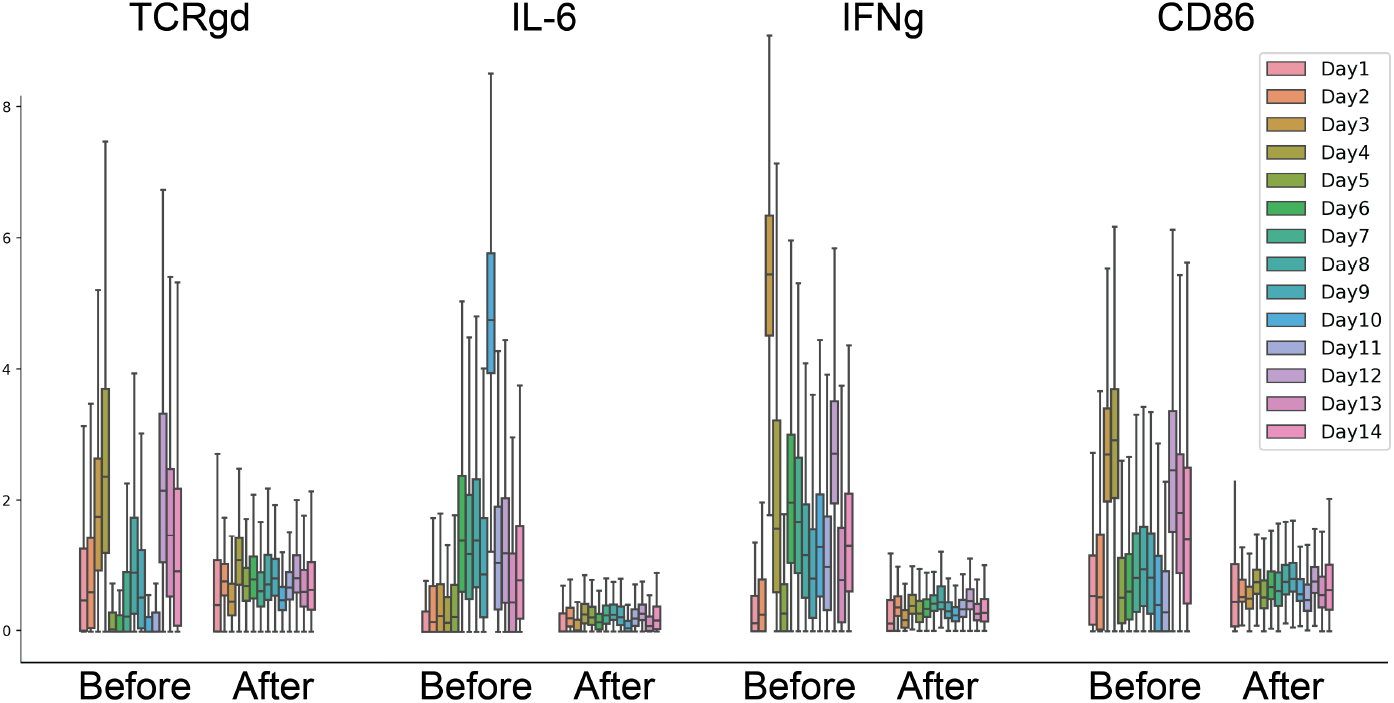
Four select marker abundances with samples grouped by day they were run on the cytometry instrument, with each day having fourteen distinct samples in the group. For each marker, the fourteen samples before batch correction are shown to the left of the same fourteen samples after batch correction.

**Supplemental Figure S8:**
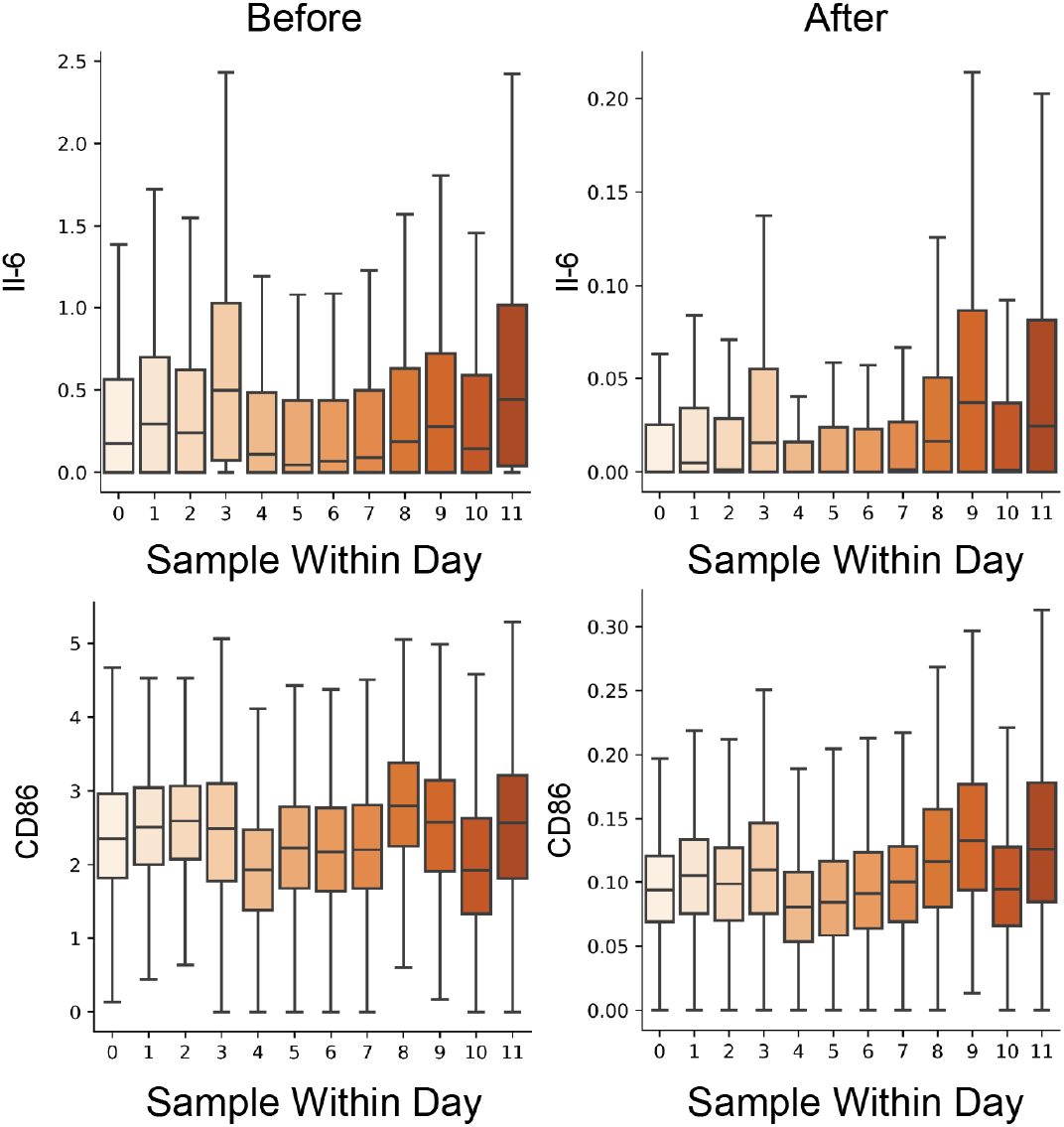
Histograms of marker expression (top: IL-6, bottom: CD86) of samples run together on the cytometry instrument on day two, separated by sample. The values for each sample and marker are shown before SAUCIE batch correction (left) and after SAUCIE batch correction (right).

**Supplemental Figure S9:**
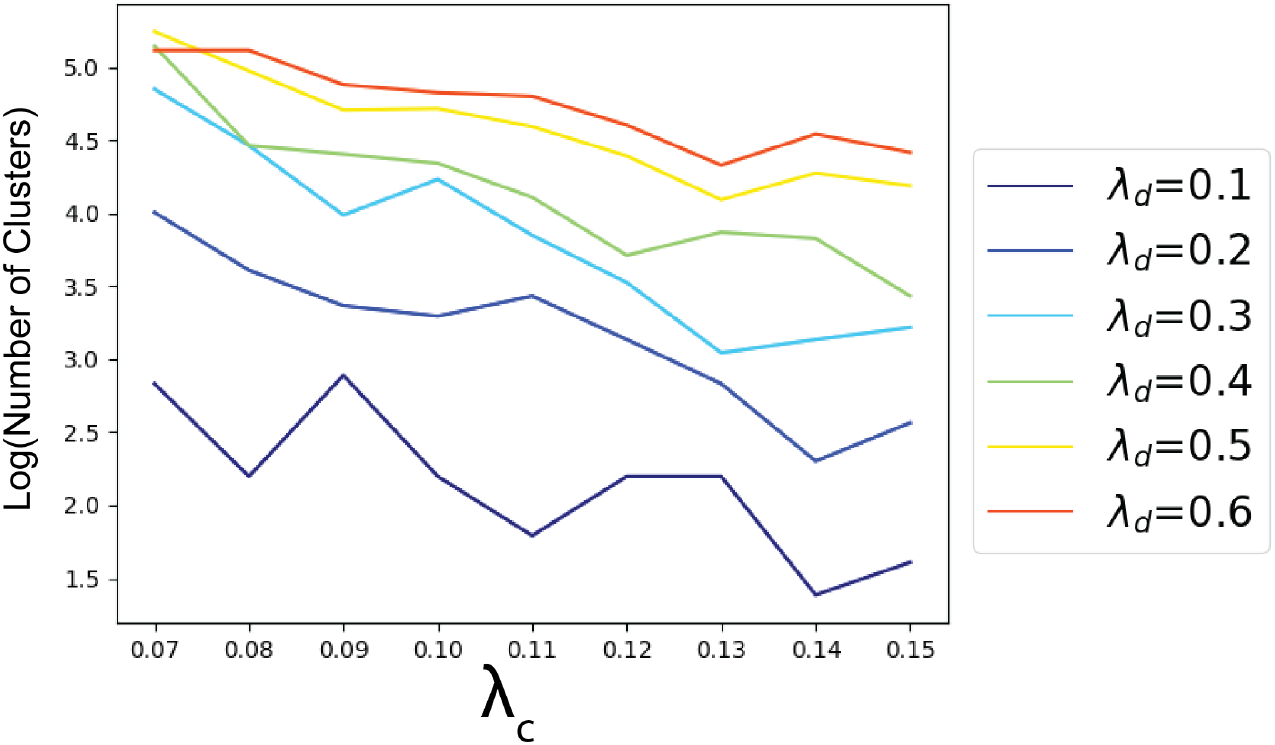
The granularity of the clustering, as measured by the total number of clusters found. Each line represents a fixed value of *λ*_*d*_ as *λ*_*c*_ increases from left to right.

**Supplemental Figure S10:**
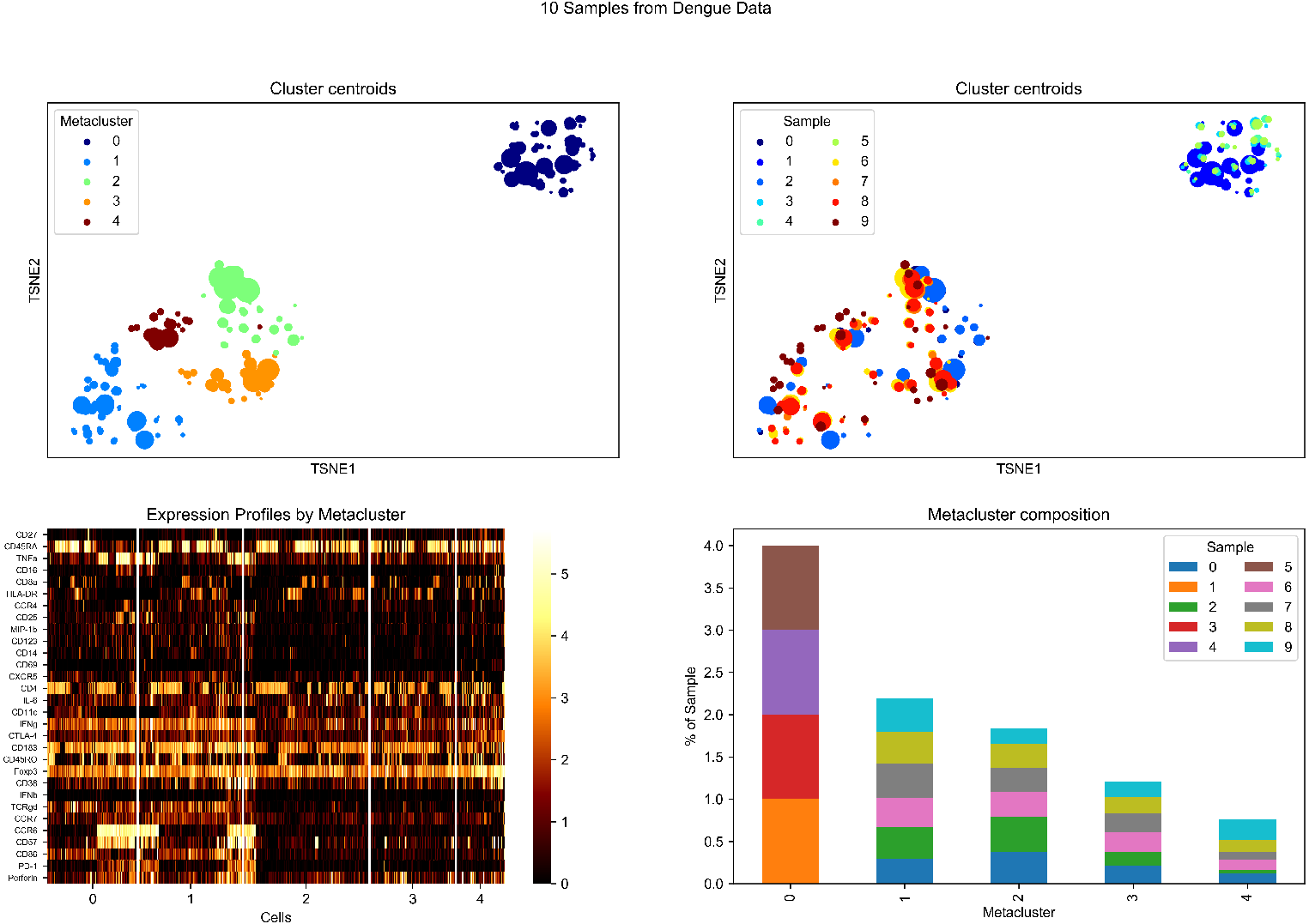
An illustration of the metaclustering process on the dengue dataset. Top left: cluster centroids embedded by tSNE and colored by metacluster, sized according to the number of cells in each cluster. Top right: cluster centroids colored by sample, also sized according to the number of cells in each cluster. Bottom left: a cell-level heatmap of expresssion grouped by metacluster. Bottom right: the composition of each metacluster by sample.

**Supplemental Figure S11:**
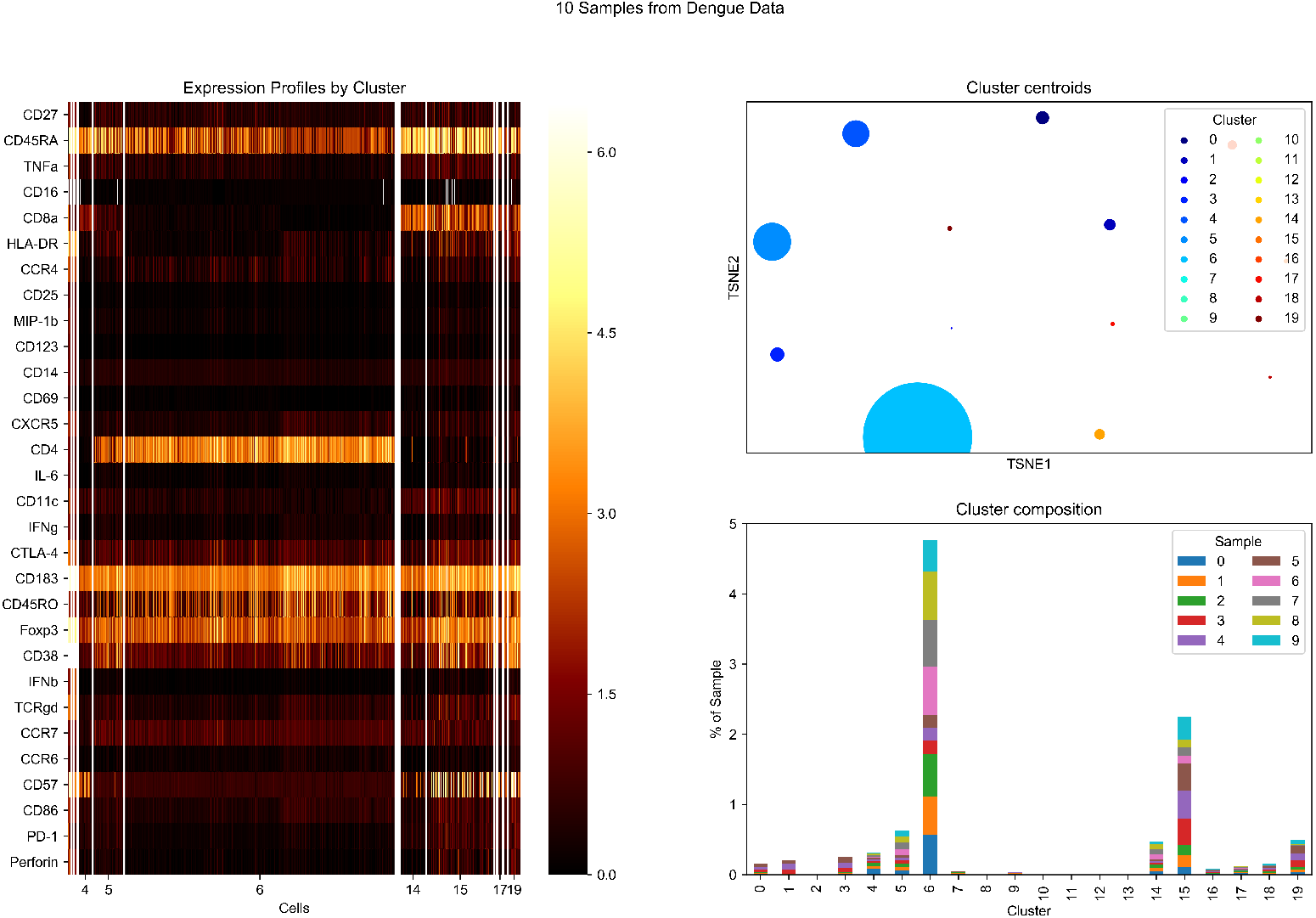
An illustration of the SAUCIE pipeline on the dengue dataset. Left: cell-level heatmap of expresssion grouped by cluster. Top right: cluster centroids embedded by tSNE, sized according to the number of cells in each cluster. Bottom right: the composition of each cluster by sample.

